# Harnessing insect olfactory neural circuits for noninvasive detection of human cancer

**DOI:** 10.1101/2022.05.24.493311

**Authors:** Alexander Farnum, Michael Parnas, Ehsanul Hoque Apu, Elyssa Cox, Noël Lefevre, Christopher H. Contag, Debajit Saha

## Abstract

There is overwhelming evidence that metabolic processes are altered in cancer cells and these changes are manifested in the volatile organic compound (VOC) composition of exhaled breath. Here, we take a novel approach of an insect olfactory neural circuit-based VOC sensor for cancer detection. We combined an *in vivo* antennae-attached insect brain with an electrophysiology platform and employed biological neural computation rules of antennal lobe circuitry for data analysis to achieve our goals. Our results demonstrate that three different human oral cancers can be robustly distinguished from each other and from a non-cancer oral cell line by analyzing individual cell culture VOC composition-evoked olfactory neural responses in the insect antennal lobe. By evaluating cancer vs. non-cancer VOC-evoked population neural responses, we show that olfactory neurons’ response-based classification of oral cancer is sensitive and reliable. Moreover, this brain-based cancer detection approach is very fast (detection time ~ 250 ms). We also demonstrate that this cancer detection technique is effective across changing chemical environments mimicking natural conditions. Our brain-based cancer detection system comprises a novel VOC sensing methodology that will spur the development of more forward engineering technologies for noninvasive detection of cancer.

## Introduction

Breath analysis is a noninvasive disease detection technique that aims to characterize the volatile chemical composition of exhaled breath, which represents the volatile chemicals present in blood and airways inside the body ^1–5^. Cancer alters cellular metabolism and these alterations are ultimately reflected in the volatile organic compound (VOC) composition of patients’ exhaled breath ^1,6,7^. It is well known that volatile cancer biomarkers are present in exhaled breath at a detectable range (parts per million to parts per trillion) ^1,8^. Recent studies have identified several putative volatile biomarkers associated with multiple cancers, including head and neck, lung, and breast ^1,8,9^. Moreover, it is presumed that cancer-induced changes in breath samples are detectable at early stages of the disease, and recent studies have supported the possibility of early cancer detection by analyzing VOCs in exhaled human breath sample ^2,5,10,11^.

Consequently, patients can potentially be screened early, noninvasively, and periodically by identifying unique exhaled breath volatile compositions indicative of cancer. Currently however, there is no gas sensing technology being used in clinical settings for cancer detection. The most commonly used volatile chemical sensing technology is gas chromatography-mass spectrometry (GC-MS), which performs individual component-wise identification of gas mixtures ^1^. Although the GC-MS-based technique is sensitive and has been shown to identify putative cancer biomarker concentrations in breath samples, it is not suitable for clinical settings, being generally slow, not portable, and requiring pre-processing and storage of samples. Moreover, this component-wise classification approach is fundamentally different from biological olfactory gas sensing and poses challenges regarding diagnostic capabilities due to internal variations of breath samples and environmental factors. Another gas sensing technology, electronic nose (e-nose) devices employ biological principles, such as combinatorial coding approaches, to achieve one shot VOC sensing ^12–14^. Although, these portable and inexpensive chemical sensors are able to process breath samples in real time, even after decades of development, they still lack specificity, cross-selectivity, and the ability to work in natural conditions ^13^.

While there are limitations in engineered chemical sensors to detect volatile compounds in natural settings reliably, biology has solved this problem over millions of years of evolution. The canine nose is the most widely used biosensor and remains the state-of-the-art approach for several gas sensing applications including homeland security and explosive detection ^15^. Trained dogs are also efficient at detecting diseases via human breath and body odors ^16–21^. However, bioassays based solely on behavior are binary, i.e., disease vs. no disease, and cannot report on different types of diseases. Insects also have an extremely sensitive sense of smell, they are easier to maintain, and can be trained behaviorally to detect specific volatiles.

In our work, we employ a forward engineering approach to ‘sniffing out cancer’ by combining a live insect brain with an electrophysiological recording platform, precision VOC delivery, and sophisticated data analysis tools. Unlike canine gas sensing which uses behavioral readouts, the olfactory neurons in the insect brain do not need to be trained to identify cancer biomarkers. Rather, cancer VOC-evoked neural response templates are used to calibrate the sensor. These VOC composition-specific neural response templates or ‘fingerprints’ can be generated for different types of diseases, which enable us to simultaneously distinguish between multiple cancers. By incorporating a live brain, this approach also harnesses the full power of a biological chemosensory array (antennae) and the associated neural computation (antennal lobe circuitry) for cancer VOC classification.

Insect olfactory sensory systems are extremely powerful and have evolved to detect low concentrations of gas molecules and minute changes in the composition of gas mixtures ^22–25^. Moreover, the locust olfactory system is well studied for odor-evoked neural coding schemes ^26–47^, and is accessible for electrophysiological recordings from multiple olfactory brain centers ^27,38,47,48^. In the insect olfactory system, VOCs are first detected by the olfactory receptor neurons (ORNs) situated in the insect antennae ^49^. Each ORN generally responds to several VOCs based on their chemical identities and concentrations ^50^. Employing a combinatorial coding scheme, 50 insect ORNs alone can detect a total of ~2^50^ chemicals, which is several trillion odorants. This enormous encoding capacity coupled with chemical specificity and specialized neural computations render the insect olfactory system extremely powerful for chemical sensing. The ORNs transmit odor-evoked electrical impulses to the antennal lobe, where the signal is processed by a complex network of excitatory projection neurons (PNs) and inhibitory local neurons. Individual PNs have broad tuning curves and they respond to several odorants and odor mixtures with varying spike rates and temporal firing motifs. The odor-evoked spatiotemporal PN population response gives rise to odor-specific neural codes, which are presumed to determine odor identity, intensity, and time course ^23,25,51^. Our previous work has identified several functional neural coding schemes that can achieve background-invariant odor recognition and novelty detection which is critical for robust detection in natural settings ^26–29^.

We obtained *in vivo* extracellular neural recordings from the PNs in the locust antennal lobe and used the neural responses for the detection of VOC mixtures emitted from human oral cancer cells. We hypothesized that locust antennal lobe PNs would respond differently to the VOC compositions associated with different oral cancer cell lines and a non-cancer cell line. We also hypothesized that this approach would be fast, reliable, and sensitive to the differences in VOC mixtures associated with different types of oral cancers. Here, we have systematically tested these hypotheses and demonstrated the feasibility and robustness of this forward engineering approach for noninvasive cancer detection.

## Results

### Cancer vs. non-cancer VOC compositions elicit distinct olfactory neuronal responses

We began by investigating odor-evoked individual projection neuron (PN) responses in the locust antennal lobe. Volatile chemical mixtures emitted from different cell cultures were delivered to the locust antenna using an olfactometer. Three oral carcinoma cell lines (Ca9-22, HSC-3, and SAS) and one non-cancer cell line (HaCaT) were grown in identical cell culture medium after seeding each cell line at the same initial cell number ^52^. All four cell cultures were grown individually in airtight flasks for 96 hours (h) to protect the emitted VOCs from contaminants. Precise amounts of the cell culture VOCs were delivered to the locust antennae for 4 seconds (s), while *in vivo* extracellular neural recordings were obtained from PNs (**Fig. 1a, b**, see Methods). Cell culture VOC samples were examined at 24 h intervals by *in vivo* PN recordings. Additionally, we used two control odorants, hexanal and undecane, which have been implicated in earlier studies as putative cancer biomarkers ^1^.

**Figure 1:**
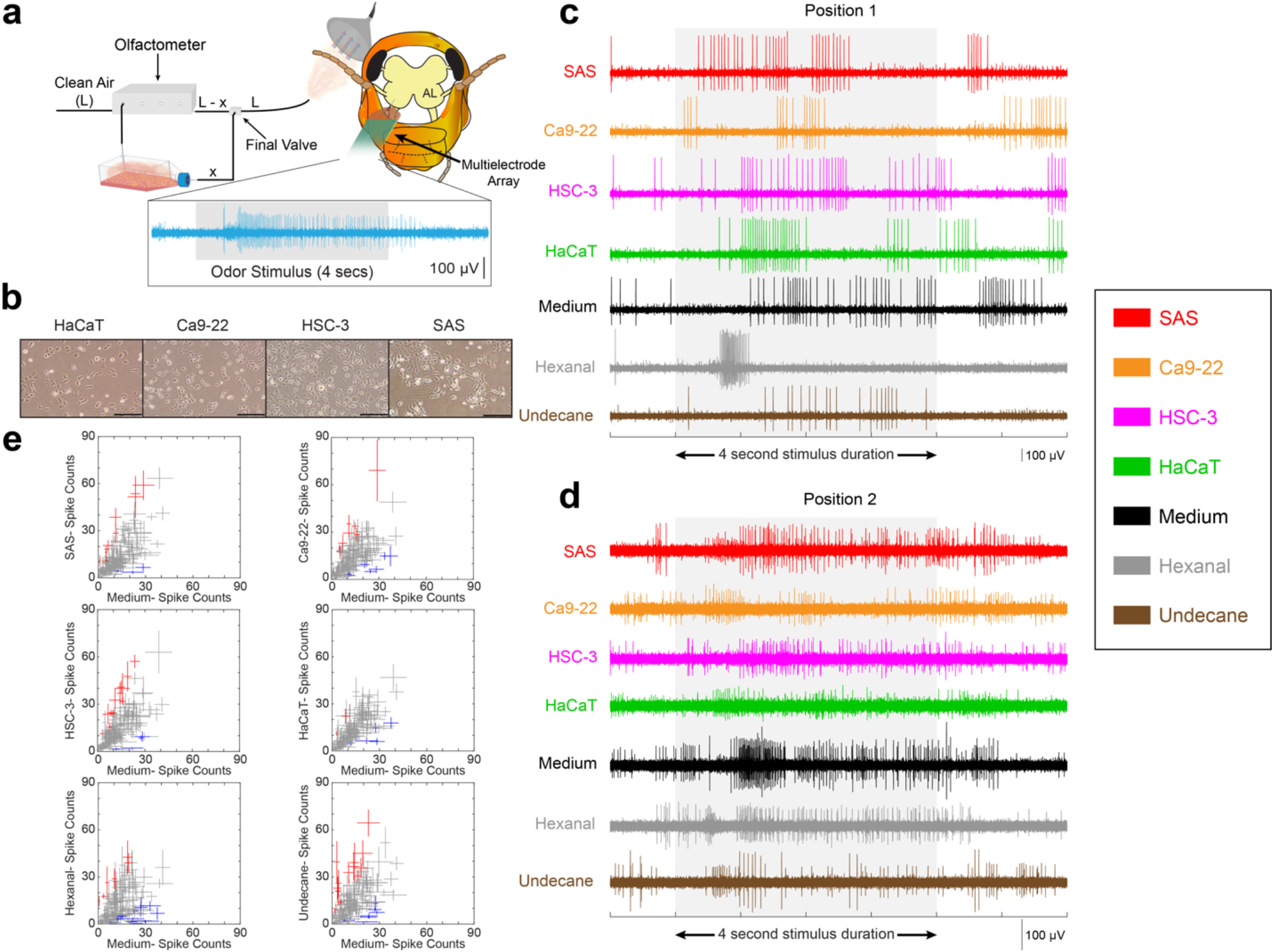
Individual projection neurons respond differentially to the oral cancer vs. control VOCs. **(a)** Schematic of the VOC delivery and *in vivo* neural recording setup. Different cell lines (cancer and non-cancer) were cultured and placed inside airtight flasks. The culture medium was the same for all cell lines. Emitted VOCs from the cell cultures were sampled periodically by injecting a fixed amount of clean air into the closed flask using an olfactometer. The duration and volume of cell culture VOCs delivered to the locust antenna were controlled by the odor delivery setup. Extracellular neural recordings were obtained from the locust antennal lobe before, during, and after odor delivery. Total airflow to the antenna was kept constant throughout the experiment and delivered VOCs were removed quickly by an exhaust placed behind the locust antenna. Bottom, a raw voltage response of a neural recording is shown for a 4 s long odor pulse. **(b)** Representative images of the four cell cultures used in the study. SAS, Ca9-22, and HSC-3 are the oral cancer cell lines while HaCaT is the non-cancer cell line. All the images are shown at 24 h post-seeding. The black scale bar indicates 200 μm. **(c)** VOC-evoked raw neural voltage responses of a recording location are shown for the three oral cancer cell lines, the non-cancer cell line, the cell culture medium, and two control VOCs (Hexanal and Undecane). The light grey box indicates the 4 s stimulus presentation window. **(d)** Similar plots are shown as in **panel c**, but for a different recording location. This location had multiple PNs, which resulted in different spike amplitudes in the multiunit voltage trace **(e)** VOC-evoked total spike counts (over 4 s) of individual PNs are compared across two stimulus conditions. For each PN, the trial-averaged total spike count is plotted with the error bars representing S.E.M. of the trial-wise variations for two stimulus conditions. All comparisons were made with the cell culture medium evoked spike counts (plotted along the X-axis). All 194 recorded PNs are plotted in every scatter plot. Individual PNs were identified after spike-sorting of the extracellular recordings. PNs that responded significantly higher (or lower) to the stimulus VOCs compared to the cell culture medium VOCs were plotted in red (or blue), respectively (P < 0.05, d.f. = 4, 28, one-way ANOVA with Bonferroni correction). PNs that did not show significant differences in total spike counts across two conditions were plotted in grey.

We observed VOC-evoked changes in neural spiking responses in most of the PNs recorded. Since PNs are broadly selective to several odor stimuli and respond to specific odorants or odor mixtures with distinctive temporal firing patterns ^22–24,51^, we targeted this neuron population for oral cancer classification. At the individual neuron level, the three oral cancer and the non-cancer VOC mixtures elicited distinct spiking responses over the odor presentation window. Raw voltage traces of representative extracellular neural recordings showed clear differentiation between the oral cancer cells, non-cancer cell, and cell culture medium. Moreover, we noted differences in PN spiking responses between the three oral cancer cell lines (**Fig. 1c, d)**.

Next, we investigated how total spike counts (over the entire 4 s stimulus window) varied for each recorded neuron corresponding to different VOC exposures (**Fig. 1e)**. To identify single neurons, spike-sorting of extracellular multi-channel recordings was performed following previously published methods ^53^. Then, we used a simple metric of VOC-evoked time-averaged and trial-averaged spike counts of individual PNs for each stimulation condition. Individual PN spike counts were summed over the 4 s stimulus presentation window and averaged across trials (*n* = 5 trials) to quantify these changes. Next, we compared the average spike count of each PN across two stimulus conditions. For example, PN spike counts corresponding to each oral cancer cell line were compared to the spike counts of the same set of PNs elicited by the culture medium VOC composition. When all recorded PNs were analyzed, several PNs showed significant changes in spike counts across two stimulus conditions (P < 0.05, d.f. = 4, 28, one-way ANOVA with Bonferroni correction). These results demonstrated that there were differences in individual PN spike counts elicited by cancer vs. non-cancer vs. control VOCs. Notice that this analysis only compared total PN spike counts corresponding to different stimuli, but differences in temporal firing motifs of individual PNs as seen in **Fig. 1c, d** were not reflected in this analysis.

### Strategies to classify oral cancer VOCs employing spatiotemporal olfactory neural responses

To incorporate the temporal spiking characteristics, we analyzed the spatiotemporal PN responses elicited by the three oral cancer cell lines, the non-cancer cell line, and the cell culture medium. To generate ‘spatial’ (neuronal identity) – ‘temporal’ (spiking dynamics) response vectors of the entire PN population, trial-averaged firing rates of each neuron were binned into 50 ms nonoverlapping time windows. Individual neuron responses were temporally aligned following stimulus onset. For this analysis, we combined spiking responses from all recorded PNs over multiple days of cell culture. This resulted in a high dimensional population neuron response, which was represented by an *n x m* matrix (**Fig. 2a**, *n* = 194 PNs; *m* = 80 time bins with 50 ms bin size over 4 s of odor presentation). Next, all recorded PN responses corresponding to each stimulus were concatenated to generate the population PN time-series data for the stimulus panel corresponding to Ca9-22, HSC-3, SAS, HaCaT, and the cell culture medium.

**Figure 2:**
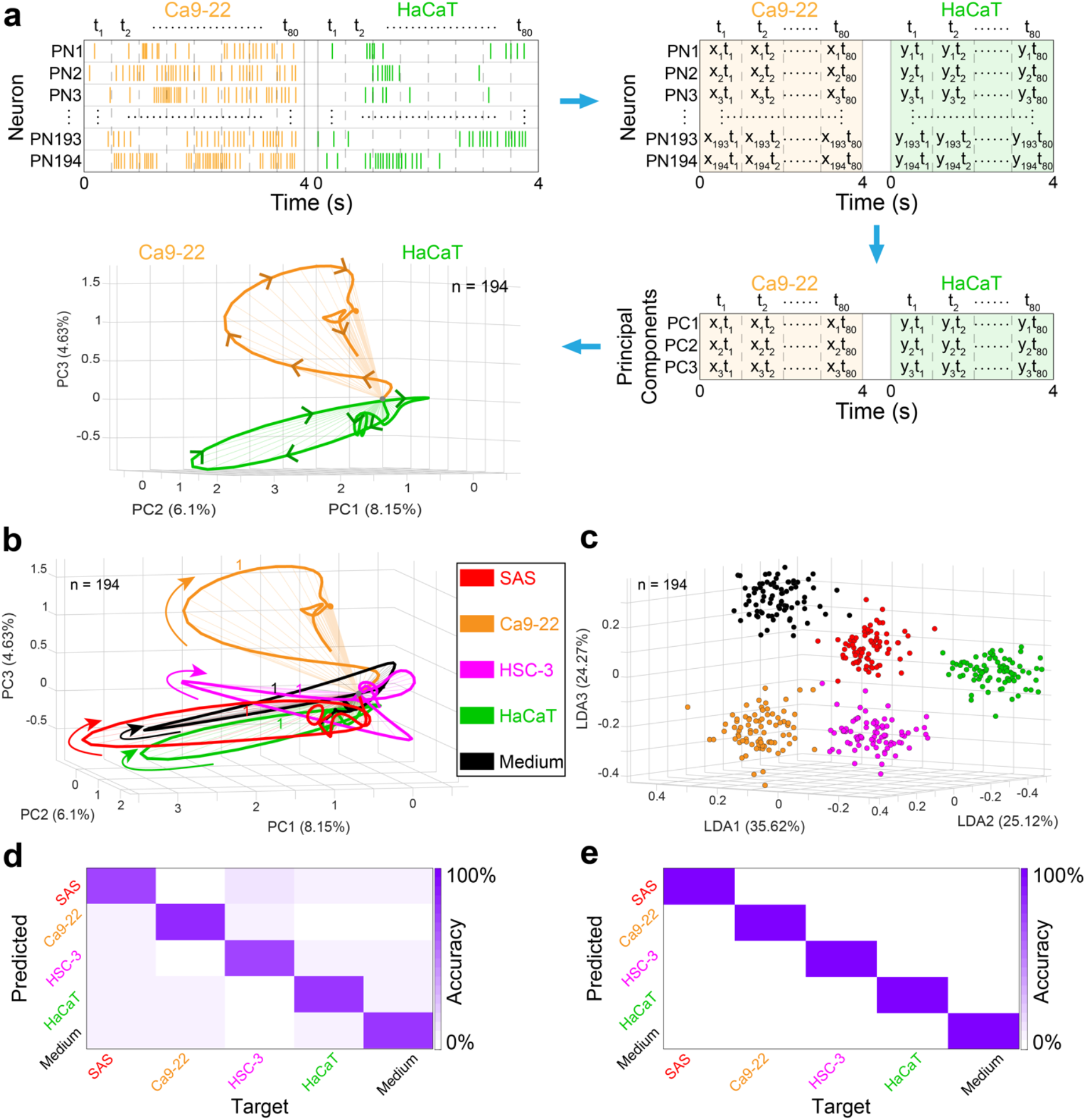
Cancer vs. non-cancer VOCs are distinguished by spatiotemporal PN responses. **(a)** Schematic representation of the spatiotemporal PN response analysis. This analysis contains spiking responses from all recorded PNs (spatial) and their response dynamics (temporal) over the 4 s stimulus presentation window. *Top left*, raster plots of all recorded PNs pooled across experiments are combined for each stimulus (represented by PN1 to PN194). Then the 4 s duration VOC-evoked spike counts are divided into 50 ms non-overlapping time bins (total 80 time bins for 4 s). Two different stimuli are used for illustration (Ca9-22 and HaCaT). Notice that the same PNs are recorded for all stimuli. Top right, a *neuron number (n=194) × time (t =80)* matrix is constructed, where each element in the matrix corresponds to the spike count of one neuron for a single time bin (denoted by *x_i_t_j_* or *y_i_t_j_*). *Bottom right*, the high dimensional population response time-series is dimensionally reduced using PCA and data corresponding to the first three principal components are kept. *Bottom left*, the 4 s time-series data is plotted along the three principal component axes. Each point is connected temporally with the next time point to generate individual VOC-evoked neural trajectories that take into account both temporal and spatial motifs of the recorded PN ensemble. Notice that the two PN trajectories corresponding to the Ca9-22 and HaCaT track along different manifolds in the principal component space. Angular separation between the two neural trajectories signifies the distinction between the two VOCs. The percentage of variance captured along the first three principal components is plotted along the axes. Because of the large number of recorded PNs and their complex response dynamics, the total variance captured along the first three principal components is low. Therefore, PCA-based neural trajectories are only used for qualitative comparisons. **(b)** Population PN trajectory plots are shown after dimensionality reduction using PCA for the three cancer cell lines (SAS, Ca9-22, and HSC-3), the non-cancer cell line (HaCaT), and the control cell culture medium. For each stimulus, PN population trajectory is plotted for 0 to 4 s of VOC exposure. Numbers along the neural trajectories indicate time in seconds from the stimulus onset. Total number of PNs used in this analysis is *n* = 194, which was computed by pooling neurons across all timepoints and replicates of the cell cultures. **(c)** Spatiotemporal PN responses (*n* =194) for the 4 s stimulus presentation window (50 ms bin size, total 80 points for each odor) are visualized after dimensionality reduction using LDA (see Methods). LDA minimizes within-class variance and maximizes the variance between classes. Numbers along the axes indicate the variance captured along that dimension. Distinct clustering of neural responses corresponding to different VOC profiles indicates that the cell culture VOCs (cancer vs. non-cancer) can be segregated based on the neural response they elicit. **(d)** Classifications of VOC-evoked population PN responses without any dimensionality reduction are analyzed by *leave-one-trial-out* cross validation analysis (see Methods). These quantitative classification results are summarized by a confusion matrix. Each column and row correspond to the target stimulus and the predicted class, respectively. Here, each 50 ms time bin of the testing trial is classified as one of the 5 target VOCs based on the minimum Euclidian distance. The high values along the diagonal of the confusion matrix indicate that most of the predicted responses match the target labels. This result signifies that information contained within the 50 ms time bins of the VOC-evoked neural response is sufficient to classify oral cancer vs. non-cancer and to distinguish different oral cancers from each other. **(e)** Similar analysis as shown in **panel d** except we classified the test trial as a whole by taking the mode of the bin-wise classification for the 4 s long trial (mode of total 80 time bins for each test trial). This trial-wise classification of VOC profiles shows flawless distinction of all 5 stimuli tested and reveals the strength of this neural response-based cancer detection approach.

To visualize these cell culture VOC-evoked spatiotemporal neural responses, we projected the high dimensional data onto three dimensions using a linear principal component analysis (PCA, **Fig. 2b**, see Methods). The points in the three-dimensional PCA subspace were connected in a temporal order to generate stimulus-specific neural response trajectories. We observed that each VOC profile generated a closed loop neural trajectory, which evolved in a unique direction. A long line of work in insect olfaction has established that the unique direction of the population PN trajectories are specific to odor identity and intensity ^23,24,26^. Our previous work demonstrated that larger angular distances between PN trajectories signify better separability between two odorants ^24,26^. Therefore, unique neural trajectories corresponding to individual VOC mixtures indicate that oral cancer VOC profiles are distinct from the non-cancer cell line. Moreover, we observed distinctions among the neural trajectories evoked by the three oral cancer cell lines (**Fig. 2b**), which signify that differences between various oral cancers can be identified by this approach as well.

To determine the separation between the cell line specific neural response clusters, we performed linear discriminant analysis (LDA) on the population PN time-series data (**Fig. 2c**). Similar to the PCA analysis, we used the population PN time-series dataset and plotted the VOC-evoked PN responses in a three-dimensional LDA subspace. This linear dimensionality reduction technique maximized the neural response cluster separation between stimuli. We observed distinct clustering of PN responses corresponding to all five stimuli, indicating that a linear classifier in a three-dimensional LDA space is sufficient to classify cancer vs. non-cancer successfully based on their corresponding VOC profiles.

To get a quantitative estimate of the classification performance, we performed a *leave-one-trial-out* cross validation analysis of the PN time-series data (see Methods, **Fig. 2d, e**). This analysis was performed on the high dimensional dataset (*n* = 194 PNs, *m* = 80 time bins) without any dimensionality reduction. First, Euclidian distances of neural response vectors at each time bin (50 ms duration) were compared between the testing and the training data (total 80 comparisons over 4 s for each test trial), generating a bin-wise classification (**Fig. 2d**). This bin-wise confusion matrix had its highest values along the diagonal, which implied a high rate of successful detection of all five stimuli. Next, we plotted a trial-wise confusion matrix by calculating the mode of the predicted responses for all 80 time bins (**Fig. 2e**). The trial-wise analysis was implemented to assign one predicted value for each test trial. This analysis showed 100% classification for all three oral cancer VOC mixtures among themselves and in comparison with the non-cancer and control VOCs. Similar dimensionality reduction and confusion matrix analyses were performed on the dataset while including the two other control odorants (**Fig. S1**). Neural trajectories corresponding to the two control odorants were significantly different from all cell culture VOCs and the confusion matrix analysis showed high classification success for all seven VOCs tested.

### Classification of cancer vs. non-cancer cells in a changing chemical background

We anticipated that emitted VOC compositions corresponding to each cell line would vary over time due to cell growth and ongoing metabolic processes in a fixed cell culture medium. We also hypothesized that the neuronal template-based VOC classification approach would be able to compensate for these variations caused by changing environments. To investigate this, the neural data that were previously combined were split and analyzed at four different time points: 24-, 48-, 72- and 96-h after seeding (**Fig. S2**). All PNs recorded at a specific time point across multiple repetitions of the cell cultures were combined to generate the population PN response vector for that time point. For example, each cell culture was repeated 7 times, and the VOC analysis at the 24-h time point resulted in a total of 42 PNs. All the cell cultures remained viable over 96-h from initiation, which was verified by manually counting healthy cells at different time points of the cell cultures (**Fig. S3, S4**).

We began by examining VOC-evoked population PN time-series data at 24-h post seeding. We noticed that dimensionally reduced neural trajectories evolved in different directions for different VOC profiles in the PCA space (**Fig. 3a**). When we performed the same analysis at 48-h, 72-h and 96-h time points, we continued to observe distinct cell line specific neural trajectories, which indicated that all the tested stimuli were distinguishable from each other at different time points of cell growth (**Fig. 3a-d**). This observation demonstrated that cultured cells started emitting VOCs specific to their identity early (~ 24-h) and remained separable over multiple days based on their emitted VOC profiles. Next, we analyzed the neural cluster separation between the three oral cancer cell lines, the non-cancer cell line, and the control medium at different time points of the cultures using LDA (**Fig. 3e-h**). Since the number of PNs recorded at each time point was low, PN response clusters showed some overlap in the LDA space. This was also reflected in the time binwise confusion matrix classification results performed in the high-dimensional space (**Fig. 3i-l**). However, the trial-wise classification result yielded 100% classification success for each test trial for all the VOCs at all four time points (**Fig. 3m-p**). Notice that we generated VOC-specific neural fingerprints at each time point of the cell cultures and performed *leave-one-trial-out* cross validation between the test trials and the training templates generated at the same time point.

**Figure 3:**
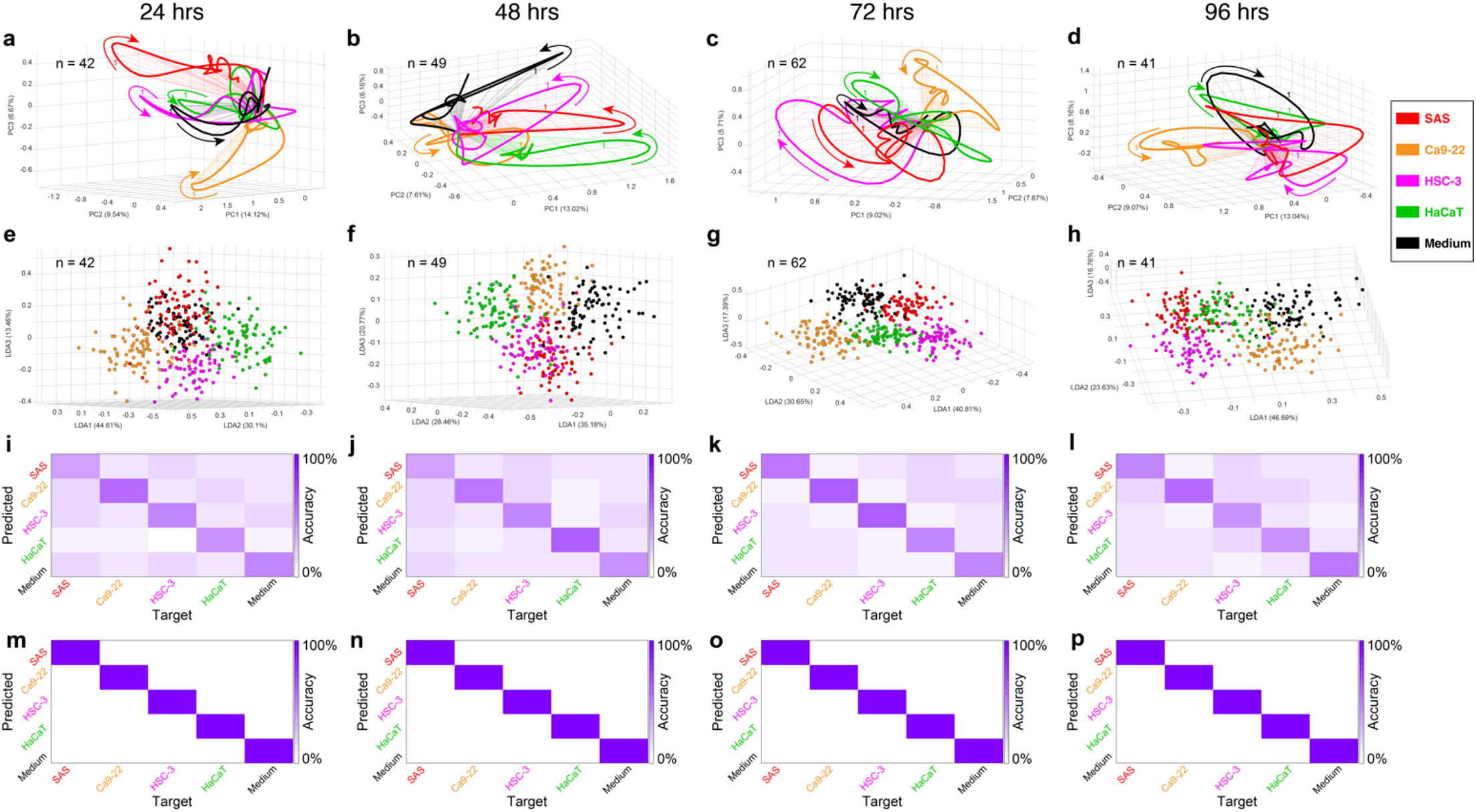
Neural response-based classification of oral cancer is robust in changing background conditions. **(a)** PN response trajectories corresponding to the cancer cell lines, the non-cancer cell line, and the cell culture medium VOC mixtures are plotted after dimensionality reduction by PCA (similar to **Figure 2**). In this analysis, neurons were pooled only across experiments performed at 24-h post seeding. A total of 42 PNs were recorded for different cell cultures at this time point. Notice that neural trajectories are distinct from each other even when the recorded neuron numbers are lower and the cultured cells are grown for only 24-h. **(b-d)** Cell culture VOC-evoked neural trajectory plots are shown for 48-, 72-, and 96-h time points of the cell cultures, respectively. At each time point, PN trajectories traversed distinct manifolds indicating the distinguishability between cell cultures over multiple days of growth. Note that different PNs (*n*) were recorded at different time points. **(e)** PN population responses cluster distinctly for different VOCs after dimensionality reduction by LDA at 24-h. **(f-h)** The same analysis as in **panel e,** used for 48-, 72-, and 96-h of cell growth. In all cases, LDA shows separability between VOC-evoked neural response clusters. **(i)** Time bin-wise high dimensional confusion matrix analysis of PN responses by *leave-one-trial-out* approach at 24-h. The confusion matrix has higher values along the diagonal, which indicates that most of the test trial time bins are classified correctly. However, the confusion matrix also has non-zero off-diagonal elements, indicative of some misclassification. **(j-l)** Similar confusion matrix plots are shown for 48-, 72-, and 96-h. **(m-p)** Trial-wise confusion matrices are shown at 24-, 48-, 72-, and 96-h of cell cultures. Here, each test trial was classified based on the mode of the bin-wise classification results. This analysis elicits diagonal confusion matrices for all cases, which indicates clear distinction of oral cancer vs. noncancer VOC profiles based on population PN spiking responses.

These results validated our hypothesis that neural response-based classification of cancer VOCs is unaffected by the variations in chemical background caused by evolution of cancer cells in the culture medium. Considering that fluctuations due to internal and external factors is a problem for current breath sensing technologies, the ability to differentiate all four cell cultures over days is a unique feat achieved by this approach.

### Neural response-based classification of cancer VOCs is fast

We investigated how short of a VOC exposure will result in robust cancer classification. We hypothesized that a neuron response-based classification approach would be fast and able to classify different VOCs with a short inter-stimulus interval (~ 1 minute). Based on the fast PN response dynamics, we anticipated that distinction between cancer VOCs would be achieved within a few hundred milliseconds of stimulus exposure. To achieve fast analyses of neural signals, we employed a different metric of neural response, which was obtained by root mean squared (R.M.S.) filtering of raw neuron voltage responses (**Fig. S5**, see Methods). Until now, all classification analyses were performed after spike-sorting of multi-unit extracellular voltage responses obtained from each recording location. However, this approach eliminated neurons that did not pass the statistical test necessary to be counted as single units. These lost signals from unresolved neurons could potentially be important for odor discrimination, therefore, we decided to employ the R.M.S.-based approach which takes into account the total energy of the signal acquired from each location. This approach was computationally less expensive, unsupervised and shown to be odor specific in our earlier work ^22^. Using the R.M.S. filtered population PN voltages, we observed distinct classification of all 7 VOCs tested (**Fig. S6**). These classification results were qualitatively similar to the results obtained from spike sorted single unit data.

To determine the speed and efficacy of this method, we performed VOC classification during four different 250 ms time segments of the 4 s stimulus presentation window (**Fig. 4**). The rationale behind choosing different time windows follows from the unique odor-evoked response dynamics of the projection neurons. PNs generally fire strongly with high spiking rates within the first ~1.5 s of stimulus onset, which is known as the ‘transient state’^23,28,32,51^. After about 2 s of stimulus exposure, the population PN firing rate converges to a stable firing rate, which stays above baseline firing but does not change significantly over the rest of the odor presentation duration. This is known as the ‘steady state’ response period. It is shown in our and others’ work that odor-evoked transient PN responses are more discriminatory^22–24^. Therefore, we expected the cell culture VOCs to display the best separation when the population PN responses are within the transient state. We observed that odor plumes took about 0.5 s to elicit spiking responses in PNs. This time corresponded to the delay between the final olfactometer valve opening and the odor plume hitting the antenna. Therefore, we chose the analysis time windows for transient PN response period as 0.5 – 0.75 s and 0.75 – 1 s and the steady state time windows as 2 – 2.25 s and 2.25 – 2.5 s (**Fig. 4**).

**Figure 4:**
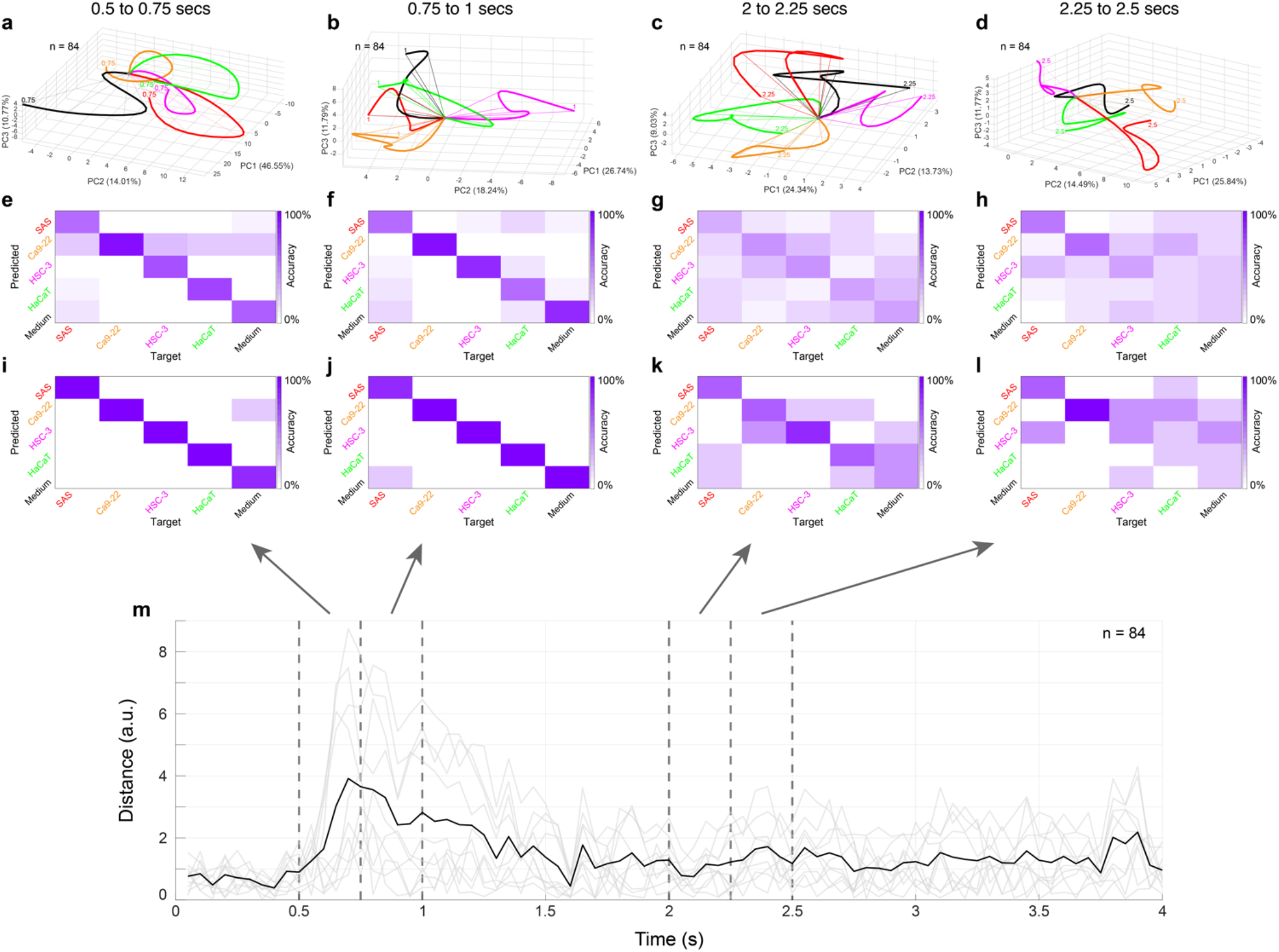
Rapid classification of oral cancer VOC profiles using neural voltage responses. **(a-d)** High dimensional neuron response vector, where each row represents R.M.S. filtered PN signals of a recording position (*n* = 84) and each column represents a 50 ms time bin (total 5 time bins over 250 ms), was dimensionally reduced using PCA and ensemble neural response trajectories are plotted (see Methods). Cancer vs. non-cancer VOC-evoked neural response trajectories are shown for the stimulus presentation windows of 0.5 – 0.7 5 s, 0.75 – 1.0 s, 2.0 – 2.25 s, and 2.25 – 2.5 s, respectively. Notice that population trajectories generated from R.M.S. filtered neural voltages are distinct within just 250 ms of odor exposures. Two 250 ms time windows are shown during the transient state of the PN response (0 to 1.5 s), while two other time windows are chosen during the steady state neural response period (2 s to the termination of the stimulus). **(e-h)** Confusion matrix analysis of the predicted vs. target responses are shown for the ensemble neural voltage time-series data for the same time windows as shown in **panel a-d**. Note that the confusion matrix analysis is done without any dimensionality reduction. Transient state time windows of 0.5 – 0.75 s and 0.75 – 1.0 s show better VOC classification compared to the steady state time windows**. (i-l)** The same confusion matrices are plotted for the trial-wise classification, which results into near perfect classification of VOCs in 250 ms time windows during transient state. Steady state windows show relatively low trial-wise classification. **(m)** Pairwise distances between ensemble R.M.S. voltages (from *n* = 84 PN recordings) corresponding to five different VOCs are plotted in light grey (total 10 pairwise distances) during the stimulus exposure (4 s). The mean of pairwise distances is plotted in black, which indicates that there are differences in ensemble PN voltage responses corresponding to different VOCs and these differences are highest during 0.5 to 1 s of the transient response period. Note that in our setup, odorants took about 500 ms to reach the antenna from the opening of the final olfactometer valve (t = 0) and therefore, the earliest transient state time window that could be chosen was 0.5 to 0.75 s.

We performed PCA dimensionality reduction analysis to visualize population neural trajectories, which showed distinct trajectories at the earliest of the time windows (0.5 to 0.75 s). The VOC-evoked neural trajectories remained distinct during both transient and steady state time epochs (**Fig. 4a-d**). Next, we performed the quantitative high dimensional confusion matrix analysis using *leave-one-trial-out* methodology. We observed better classification during transient state time windows compared to the steady state time windows, evident from the higher value of diagonal elements in the confusion matrix shown in **Fig. 4e, f** in comparison to **Fig. 4g, h**. Trialwise classification also showed better predictability during transient state response periods (0.5 – 0.75 s and 0.75 – 1 s) compared to the steady state segments (2 – 2.25 s and 2.25 – 2.5 s, **Fig. 4i-l**). Finally, when we compared the pairwise R.M.S. response distances of the PN population elicited by all 5 VOCs, we observed the largest separation was also during the transient periods. These sets of results demonstrated that neural response-based cancer classification is fast and only requires 250 ms of neural data from stimulus onset to distinguish oral cancers from controls.

To verify that a one-minute inter-stimulus interval is sufficient for the VOC classification and our results are consistent with the PN response dynamics, we employed the R.M.S.-based classification analysis on the baseline, transient, and steady state epochs of the population PN response (**Fig. S7**). Each analysis epoch was 1.5 s in duration and the 0.5 s delay for the odor stimulus to reach the antenna was included in the pre-stimulus period. We observed no classification in the baseline period (−1 to 0.5 s), but VOC classification was distinct during the transient (0.5 to 2 s), and steady state (2 to 3.5 s) periods. Overall, VOC-evoked neural responses during the transient period yielded best classification results as expected from the PN response dynamics ^24^.

## Discussion

Current state-of-the-art gas sensing technologies (e.g., GC-MS) analyze a gas mixture to identify individual chemical components and their concentrations. GC-MS has shown promise as a diagnostic technology for a number of diseases including asthma, COPD, cystic fibrosis, diabetes, and cancer ^11,54–60^. However, there does not exist a single VOC biomarker that is indicative of a specific type of cancer. Instead, subtle changes in VOC compositions indicate altered metabolic processes corresponding to a particular cancer. Moreover, chemicals such as nitric oxide, nitrogen dioxide, and ethane have been observed to be key biomarkers in a number of diseases, yet are difficult to detect with GC-MS due to the lag in sampling-to-processing time ^61^. This component wise VOC mixture classification approach is also hindered by the variability in VOC compositions in exhaled breath between individuals (internal factors) as well as due to the presence of any background odorants (external factors) in the environment.

While GC-MS has proven essential for volatile chemical identification, the desire for point-of-care clinical implementation has fueled interest in developing low-cost, portable sensors ^12,13^. Electronic noses have increased in popularity owing to advancements in materials science, nanotechnology, and pattern recognition algorithms. These devices have demonstrated the ability to distinguish between the ‘breath prints’ of healthy controls and those afflicted with diseases, such as cystic fibrosis ^62^, cancer ^63–65^ and others ^65–69^. These point-of-care sensors lack GC-MS’ exceptional chemical sensitivity, typically only achieving detection thresholds in the parts-per-million and parts-per-billion ranges. Although electronic noses have shown promise in some disease classification, sensitivity is of concern when considering clinical implementation, as endogenous volatile compounds are typically found on the order of parts per billion to parts per trillion ranges in exhaled breath ^70,71^. Moreover, differences in environmental volatiles between patient sampling can lead engineered sensors to produce false classifications and diagnoses ^72^. Therefore, a background-invariant chemical sensing system is essential for the evolution of breath-based clinical diagnostics.

In biological olfaction, natural selection has forced animals to develop highly sensitive olfactory capabilities while preserving chemical specificity. In the olfactory sensory system, a target VOC mixture as a whole is encoded by a distinct neuronal response template (or a neuronal ‘fingerprint’ of a VOC), while a different gas mixture is uniquely encoded by a different neuronal fingerprint. It is important to note that biology does not perform component-wise classification of gas mixtures, but instead achieves optimal separation between the VOC-evoked neural fingerprints. Through experience, biological systems assign meaning to those neuronal fingerprints (e.g., food vs. harmful odors). For example, the implementation of a neuronal template-based classification approach enables honeybees to detect minute changes in odor mixtures, such as differentiating between a flower with nectar vs. the same flower without nectar based on smell alone. We utilized these biological neural coding schemes in the brain-based cancer detection approach. Based on the odor-specific population neuron responses, we first constructed neuronal fingerprints for the target VOC mixtures (training templates, **Fig. 5**). While testing an unknown VOC sample, we recorded the responses of the neuronal population and determined how well that test template matched the pre-established training templates (e.g., by measuring Euclidian distance). Based on the best match between the training and testing templates, we determined the identity of the unknown odor (**Fig. 5**). Note that the performance of this approach does not depend on neuronal identity, which will vary across experiments. This approach is based on recording from a large neuronal population (~40-50 PNs) and finding distinct training templates using known volatiles (calibration process) and then testing unknown VOCs to determine their identity using biological neural computational schemes.

**Figure 5:**
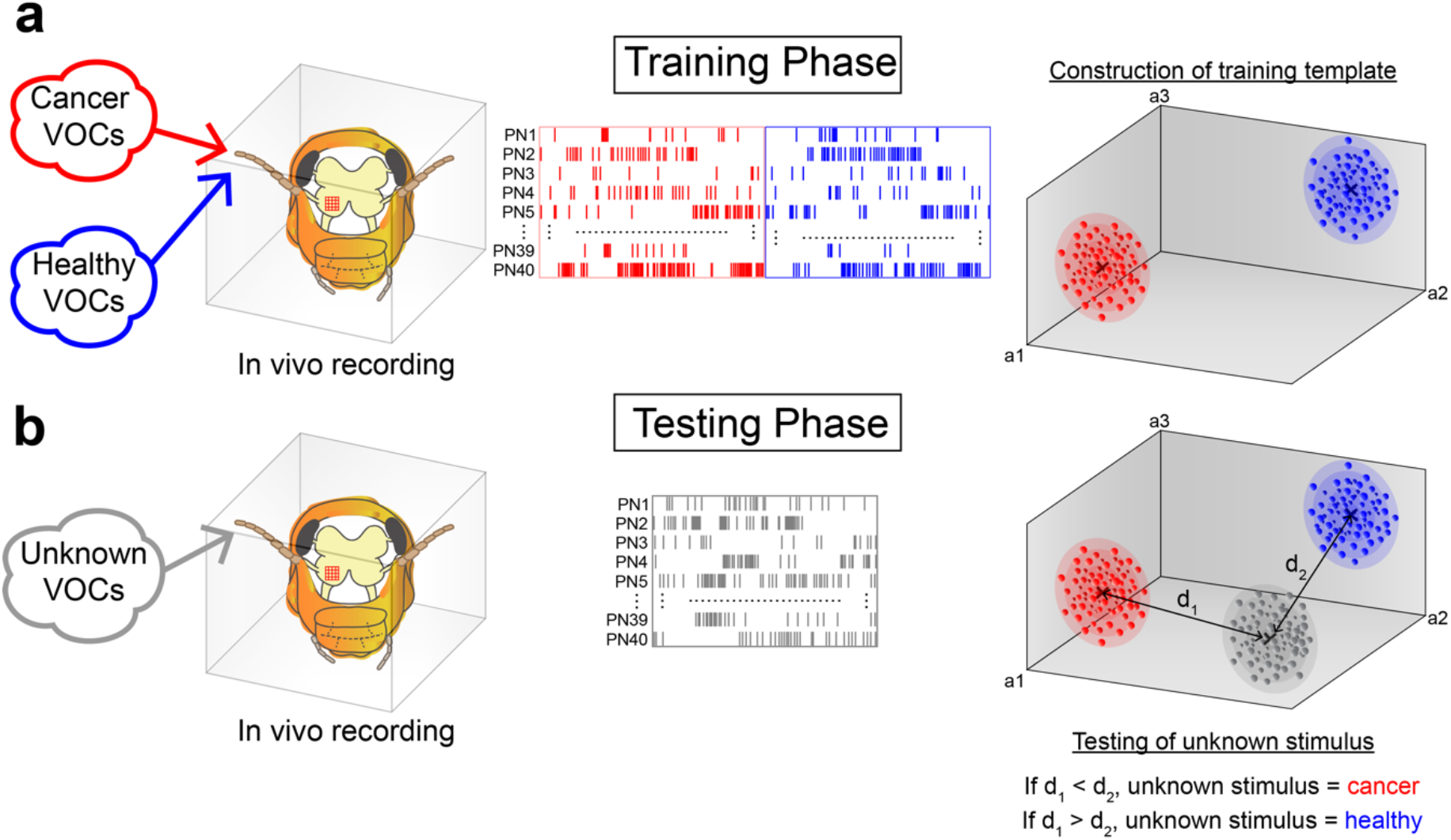
Principles of neuronal response-based noninvasive cancer detection. **(a)** This approach employs VOC mixture-evoked neural response templates for distinction between cancer and healthy samples. During the training phase, target (e.g., breath samples of oral cancer patients) and control (e.g., healthy human breath samples) VOC mixtures will be exposed to the locust antenna and *in vivo* neural recordings will be obtained from multiple projection neurons in the antennal lobe, simultaneously. The entire *in vivo* electrophysiological setup will be placed inside a closed Faraday chamber with inlet and outlet port for VOC delivery and removal. Trained VOC-evoked population neural responses will be used to construct optimally separated heathy vs. cancer clusters in the neural space as illustrated in our analysis. Our results indicate that ~40 recorded PNs is sufficient for classification of multiple oral cancer cell lines from healthy controls. Notice that the training/calibration will be performed for each brain-based sensor, where the separation between target VOCs will be maximized by optimal placement of the microelectrode array in the antennal lobe. **(b)** During testing phase, unknown VOC samples (e.g., breath sample of an early-stage oral cancer patient) will be presented to the antennae and neural responses will be obtained from the same set of neurons. In our study, we have used the minimum Euclidian distance between the unknown sample and the healthy vs. oral cancer neural clusters as the classification metric. However, other distance metrics can be used to classify unknown VOCs. Since the PN responses reach near baseline within 2 s of odor onset and we have demonstrated that reliable classification can be performed for one minute inter stimulus interval, this technique can work as a high throughput cancer screening device.

There is overwhelming evidence that cell metabolism is altered in cancer cells relative to normal cells as they switch from glycolysis to oxidative phosphorylation (OXPHOS) for energy production leading to changes in VOC compositions that are vented in exhaled breath of patients ^1,6,7,73^. We have shown that the cancer cell lines in this study demonstrated functional increases in both glycolysis and OXPHOS relative to the non-cancer cell line ^52^. Such metabolic profiles are only now being characterized in cancer biology, and a similar coincident increase in glycolysis and OXPHOS has only been documented in a resistant clone of PDAC cancer stem cells (CSC) where suppression of the c-myc oncogene and an increase in peroxisome proliferator-activated receptor-gamma coactivator-1 alpha (PGC-1a) underlie both OXPHOS and glycolysis ^74^. These metabolic differences account, in part, for the variations in VOCs from these cells. We observed that all the cell lines were healthy throughout the time of culture, however, cell proliferation rates were not optimal (Fig. S4). In order to retain maximal VOC-populated headspace, the cells were grown in a closed flask and without changing the culture medium. HEPES buffer was added to the medium to maintain the optimal pH in the airtight culture. Use of one medium for all cell lines was essential for comparison to the medium only control but may have impacted how the various cell lines grew in culture.

The concept of using canines as a way to detect disease began in 1989 when a dog alerted its owner to a suspicious mole that turned out to be cancer ^75^. Since then, canines have participated in a plethora of studies in cancer detection, including continued studies for melanoma ^20^, bladder and prostate cancer via urine ^17,76–78^, ovarian carcinomas from both blood and *ex vivo* samples ^79^, and breast and lung cancers from exhaled human breath ^16,19^. Canines have also been used to detect other types of sickness, such as hypoglycemia in patients with type I diabetes ^80^, identify stool samples containing *Clostridium difficile* from patients admitted to a hospital ^81^, and even as a diagnostic tool for covid-19, correctly identifying infected patients from patients’ clothing, masks, or breath samples ^82^. The successes in disease detection by canines led researchers to look into other animals, such as the African giant pouched rat, which has been used successfully to identify tuberculosis in human sputum samples ^83,84^. Another alternative has been to use insects. Researchers have successfully used the honeybee proboscis extension reflex (PER) to detect both covid-19 from mink throat swabs ^85^ and tuberculosis odor biomarkers ^86^. Cells lines of human breast and lung cancers have also been distinguished from healthy tissue by both fruit flies and _ants_ ^87,88^.

However, all these studies used behavioral training of animals which had a limited binary output and can be impacted by the animals’ innate behavioral preferences. Until now biologically inspired VOC detection efforts have mainly been directed towards reverse-engineering the biological olfactory system’s functionality and implementing those rules in e-nose devices. Some research groups have integrated a few live olfactory sensory receptors in engineered platforms ^89^, but those devices still lack in chemical discriminability and long-term performance. For example, e-nose devices have recently incorporated live biological olfactory receptors as chemosensor’s ^14,90–94^, including insect olfactory receptors ^95–99^. However, the integration of biological sensors into engineering platforms has proven challenging ^13^. Overall, it has become evident that it will be challenging to reverse-engineer highly efficient and intricate biological olfactory sensory systems for diagnostic means anytime soon, a skill which biology has perfected over millions of years of evolution.

Here, we took a forward engineering approach by ‘hijacking’ an insect brain to detect oral cancers from their VOC signatures. We combined *in vivo*, multi-electrode, population neuronal recordings with a multi-channel micro-amplifier, high speed data acquisition, and biological neural computations to achieve noninvasive cancer detection. This approach is fundamentally different from current gas sensing devices and animal behavior-based disease detection as it uses a fully functional biological chemosensory array (antennae) and olfactory neural circuits as a gas sensor, and neuronal ‘fingerprints’ of cancer VOC profiles as decoding schemes. We envision this study as the first step in ‘sniffing out cancer by the insect brain’ research that we can employ to detect cancer from human breath. Here, we have performed neural recordings from the brain of live animals whose chemosensory array (antennae) were exposed to VOC mixtures produced by cancer cells in culture. This *in vivo* neural recording technique can be portable as shown in our previous work ^22^. In the future, we plan to employ an antennae-attached-whole-brain (without body) in a portable and closed chamber that prolongs brain viability. This cyborg VOC sensing device will be ideal for real-time analysis of breath samples while its rapid detection ability will promote high throughput screening of a large number of VOC samples. Our next objective will be to increase the neural recording capacity and extend the brain viability for several days in a closed chamber as we progress towards the development of a portable, one-shot, point-of-care brain-based VOC sensor.

## Methods

### Electrophysiology experiments

All neural recordings were conducted on post-fifth instar locusts (*Schistocerca americana*) of either sex raised in a crowded colony. For *in vivo* extracellular recordings, locusts were immobilized on a surgical platform and antennae were stabilized. Surgery was conducted following a previously published method^24,48^. Briefly, a batik wax bowl was constructed to isolate the head region and subsequently filled with a room temperature, physiologically balanced locust saline solution. Exoskeleton and glandular tissue were removed until the brain was fully apparent, and the antennal lobes were desheathed following treatment with protease. A commercial Neuronexus 16-channel silicon probe (A2×2-tet-3mm-150-150-121) with impedances between 200-300 kΩ was used for PN recordings. Voltage signals from PNs were recorded by inserting electrodes about 100 μm into the antennal lobe. A silver-chloride ground wire was placed in the saline bath. Voltage signals were sampled at 20 kHz and digitized via an Intan pre-amplifier board. The digital signals were transmitted to a recording controller and successively visualized and stored using the Intan graphical user interface.

### Odor stimulation

A commercial olfactometer (*Aurora Scientific, 220A*) was used for precision odor stimulus delivery. Purified, zero-contaminant air was used as the carrier stream. Throughout the entirety of the experiment, a constant 200 sccm air flow was passed to the locust antenna via a 1/16 in. diameter PTFE tube, positioned approximately 2-3 cm from the last antennal segment.

During stimulus exposure, 40% (80 sccm) of the carrier stream clean air was replaced with the cell culture VOCs or other odorants. To eliminate any neural response due to sudden changes in airflow during odor delivery and removal, we kept the total volume of the airflow constant before, during, and after odor delivery. Stimulus duration for all experiments was 4 s. Each stimulus was repeated five times with an interstimulus interval of one minute. A 6” diameter funnel pulling a slight vacuum was placed immediately behind the locust antennae to ensure swift removal of odorants. The order of odor stimuli was pseudorandomized for each experiment.

### Cell culture

Human oral squamous cell carcinoma (OSCC) cell lines derived from the gingiva (Ca9-22), tongue (SAS), and a site of lymph node metastasis in tongue (HSC-3) were obtained from the Human Science Research Resources Bank (Osaka, Japan). The immortalized normal human epidermal keratinocyte cell line (HaCaT) was obtained from Cell Lines Service (Eppelheim, Germany). As a non-cancer control, HaCaT cells were chosen because they are a nontransformed, immortalized, non-tumorigenic cell line and are widely used to mimic normal stratified squamous epithelium of the oral mucosa^100,101^. The cells were all seeded at a density of 1 x 10^6^ in T-25 flasks (Nunc™ EasYFlask™ 156340, Thermo Fischer Scientific, MA, USA) with airtight caps. Airtight T25 flasks were constructed prior to the start of any experiment. Inlet and outlet 19-gauge needles were inserted into each flask and stabilized using a low-volatile, two-part epoxy at least 24-h prior to cell seeding. All the cells were cultured at 37°C in 5% CO2 using 5 mL of Dulbecco modified Eagle medium (DMEM, Thermo Fisher Scientific, MA, USA)–high-glucose (4500 mg of D-glucose/liter) medium with 25 mM HEPES and supplemented with 10% fetal bovine serum (FBS) (Biowest, France), and 1% penicillin/streptomycin (Thermo Fisher Scientific, MA, USA). HEPES was used for maintaining the pH values of the cell culture medium. Cells were allowed to grow for four consecutive days and electrophysiological data were collected at each 24-h timepoint post-seeding. Seven replicates of the four-day experiment were conducted. Flasks were maintained with a regulated temperature of 37°C and only removed while conducting experiments (less than 10 minutes). Five mL of the same cell culture medium was also placed in an identical T25 flask and kept in the same conditions as the cell cultures. Hexanal and undecane (1 % v/v in 5 mL mineral oil) were kept in identical T25 flasks and maintained at 37°C during experiments.

### Cell culture imaging and cell counting

Prior to each electrophysiology experiment, cell cultures were imaged using an optical microscope (Olympus CKX53). A total of ten images were taken at different pseudorandom locations throughout each flask. Cells were manually counted from every image (n = 1120 total images) using FIJI/ImageJ. The images from each flask taken at each 24-h timepoint were averaged and then converted to the total cell count in each flask. Mean and standard error of the mean (S.E.M.) were calculated for the total cell counts for each timepoint across seven replicates. One-way ANOVA with Bonferroni correction due to multiple comparisons was then used to determine if the cell count at each 24-h timepoint had statistically significant differences (P < 0.05, d.f. = 6, 16, one-way ANOVA with Bonferroni correction).

### Data analyses

Data was imported into MATLAB and high pass filtered using a Butterworth filter to remove any frequency components below 300 Hz. The data was analyzed by custom-written code in MATLAB.

### Spike sorting

For spike sorting analysis, all data was processed with Igor Pro using previously described methods ^53^. Detection thresholds for spiking events were between 2.5-3.5 standard deviation (SD) of baseline fluctuations. Single PNs were identified if they passed the following criteria: cluster separation > 5 SD, inter-spike intervals (ISI) < 10%, and spike waveform variance < 10%. A total of 194 PNs were identified using spike sorting from 23 locusts.

### Scatter plots

The total number of spikes for each PN during the four s of odor stimulus presentation was computed for each trial (total 5 trials of each stimulus). The mean spike counts ± S.E.M. across trials for each PN were then plotted for two stimulus conditions along X- and Y-axes (e.g., SAS spike counts vs. culture medium spike counts). One-way ANOVA with Bonferroni correction due to multiple comparisons was then used to determine if each neuron had statistically significant differences in mean spike counts to different conditions (P < 0.05, d.f. = 4, 28, one-way ANOVA with Bonferroni correction). Neurons with a statistically significant increase/decrease in spikes along the vertical axis compared to the horizontal axis were plotted in red/blue, respectively. Statistically nonsignificant differences were plotted in grey (**Fig. 1e**).

### R.M.S. transformation of PN voltage response

The filtered data was trimmed to the time window of interest. All data were passed through a 500-point continuous moving R.M.S. filter followed by a smoothing step via a 500-point continuous moving average filter. Stimulus-specific baseline values were calculated as the average voltage over all time bins for the two s prior to stimulus onset. Baseline responses were averaged over all trials and subsequently subtracted from the data to obtain the ΔR.M.S. values. These values were then binned according to the specified bin size and the average of each bin was computed. For each recording location, R.M.S. transformed voltage data of each tetrode were averaged together (**Fig. 4**).

### Dimensionality reduction analyses

We performed two methods of dimensionality reduction – Principal Component Analysis (PCA) and Linear Discriminant Analysis (LDA). In PCA, we binned baseline subtracted, spike sorted PN signals in 50 ms non-overlapping time bins and averaged over trials (n = 5, each stimulus was repeated 5 times with a 1 min inter-stimulus interval). The baseline response was calculated for each PN by averaging the firing rate over 2 s time windows immediately before stimulus presentation across trials. Recorded PNs were pooled across multiple experiments. For example, in **Fig. 2**, spike sorted and binned responses of all recorded PNs (194 total) over 24- to 96-h of cell culture were combined to generate a *PN number (n=194) × time (t =80)* matrix, where each element in the matrix corresponds to the spike count of one PN in one 50 ms time bin. Similar PN population time-series data matrices were generated for each stimulus. PCA dimensionality reduction analysis was performed on the time-series data involving 5 odorants (SAS, Ca9-22, and HSC-3, HaCaT, and culture medium) and directions of maximum variance were found **(Fig. 2)**. The resultant high-dimensional vector in each time bin was projected along the eigenvectors of the covariance matrix. Only the three dimensions with the highest eigenvalues were considered for visualization purposes and data points in adjacent time bins were connected to generate low-dimensional neural trajectories. The trajectories were smoothed using a third order IIR Butterworth filter (Half Power Frequency = 0.15). Finally, all trajectories were shifted to begin at the origin to examine stimulus-specific response dynamics and trajectory divergence. A similar approach was used for PCA analysis in **Fig 3,** except recorded PNs were separated based on the cell culture time points (e.g., 24-, 48-, 72-, and 96-h) and PCA analysis was done separately for each cell culture time point. For LDA analysis, the same population PN time-series data matrix was used. Here, we maximized the separation between interclass distances while minimizing the within class distances. To visualize the data, time bins were plotted as unique points in this transformed LDA space and stimulus-specific VOC clusters became readily apparent (**Fig. 2, 3**). The same PCA and LDA analyses were applied to the baseline subtracted R.M.S transformed population PN time-series data (**Fig. 4**).

### Classification analysis

To obtain a quantitative estimate of classification performance, we performed a *leave-one-trial-out* cross validation method. During each iteration, population PN time-series data from one trial was used as the test data and the remaining trials were used to train a linear classifier (total 5 trials for each stimulus). The linear algorithm aimed to generate a model based on the training data set within the original high-dimensional encoding state space to effectively classify testing data. By considering time bins as points in a high-dimensional space, we were able to calculate an average response vector for the training data corresponding to each stimulus. These neural templates were then used to classify individual time bins of each testing data set. The minimal norm distances between each point corresponding to a time bin of the testing data and the previously calculated average response vectors for the training data were used to assign class identities. The Euclidean (L^2^) norm was used to quantify classifier predictability in most cases. For results involving R.M.S.-transformed data, the Manhattan (L^1^) norm was selected as it outperformed the Euclidean norm metric in terms of classifier prediction accuracy (**Fig. 4 and S6**)._Furthermore, a winner-take-all approach, was also incorporated to calculate the most likely predicted class for each trial. This was performed by considering the mode of all predicted time bins as the trial-wise class identifier. Model performance was illustrated using a confusion matrix, which compared the predicted responses to the true class labels. A fully diagonal matrix indicates 100% classification accuracy.

## Supplementary figure captions

**Figure S1:**
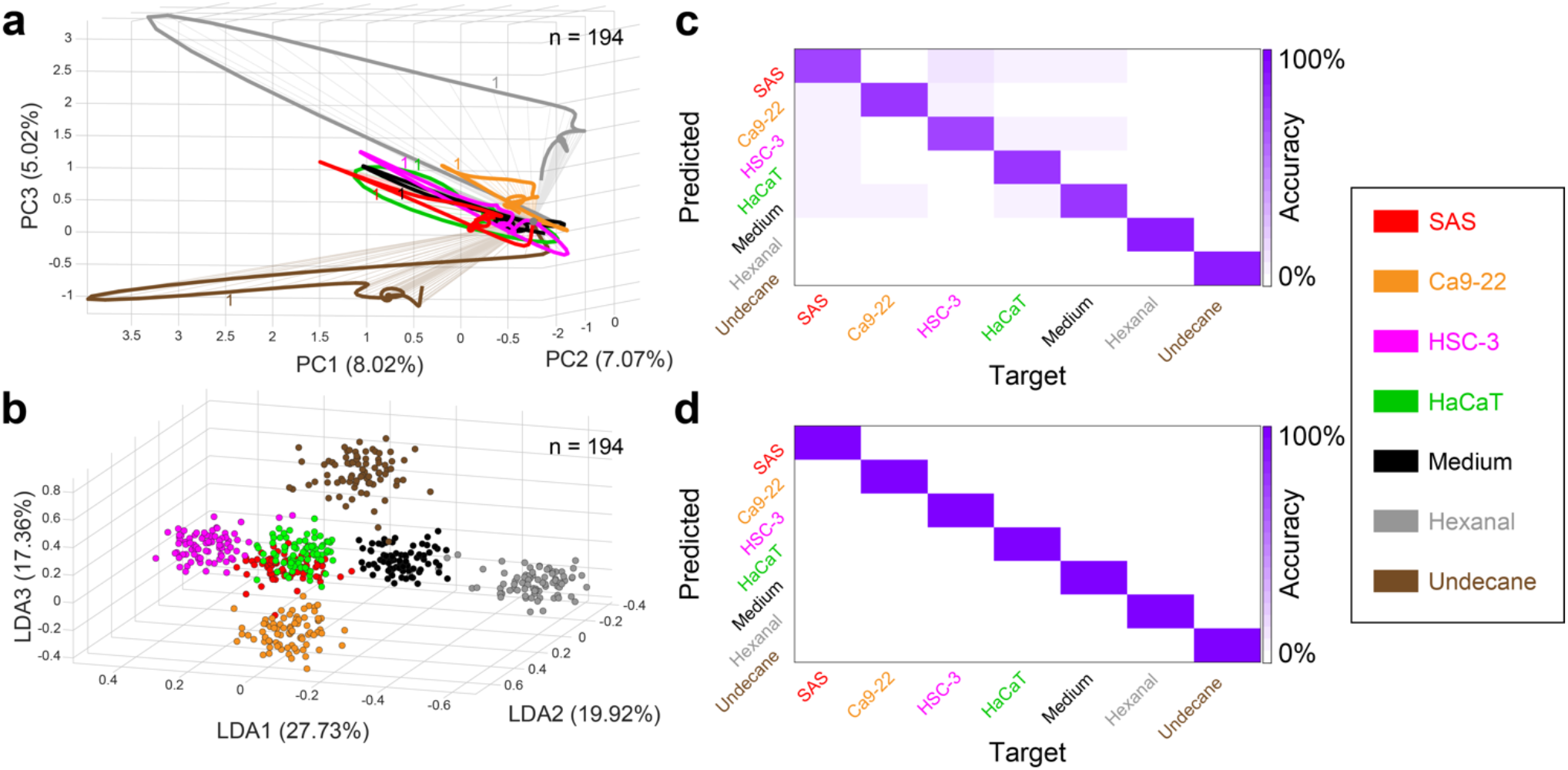
PN population response-based classification of the entire stimulus panel. **(a)** Ensemble neural trajectories over the 4 s stimulus presentation window are shown for all 7 stimuli after PCA dimensionality reduction. Volatiles from putative cancer biomarkers (Hexanal and Undecane 1% v/v diluted in mineral oil) elicited PN responses that traced different manifolds than those from the cell lines and control media. Numbers along trajectories indicate time in seconds from the stimulus onset. Total number of PNs used in this analysis is *n* = 194. **(b)** Population PN responses corresponding to each stimulus plotted in 3-dimensional LDA space also showed separability between response clusters. **(c)** Quantitative classification was performed using a *leave-one-trial-out* cross-validation methodology to train and test a linear classifier in the highdimensional feature space. The time bin-wise confusion matrix shows highest values along diagonal for all cases which indicates successful classification of all 7 stimuli using PN time-series data. **(d)** Trial-wise confusion matrix is plotted for all stimuli.

**Figure S2:**
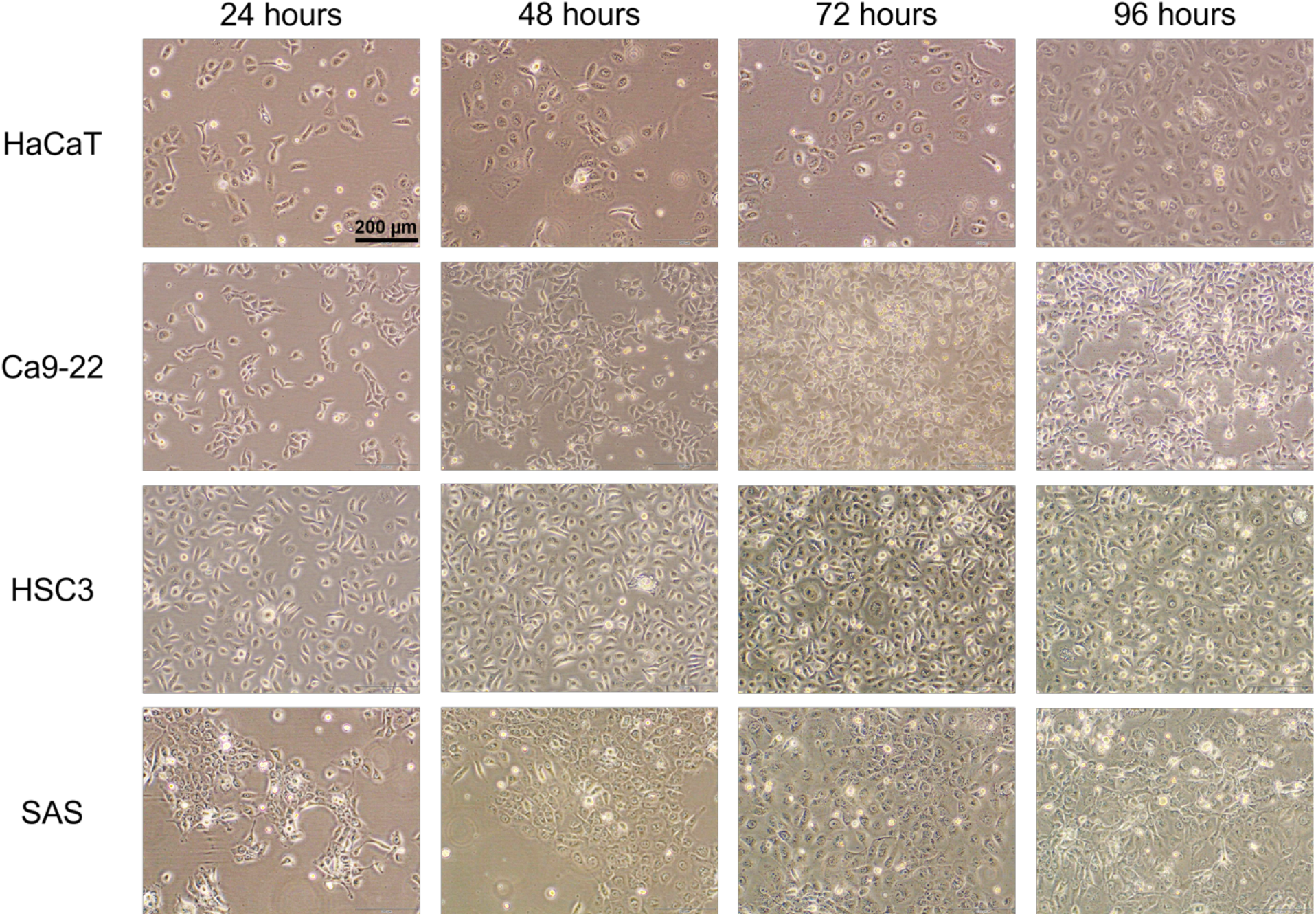
Representative images of cell cultures over days. Images are shown for a replicate of the cell culture used for electrophysiological recordings. All four cell lines are shown over four days. Healthy cells were observed at all four time points (24-, 48-, 72-, and 96-h).

**Figure S3:**
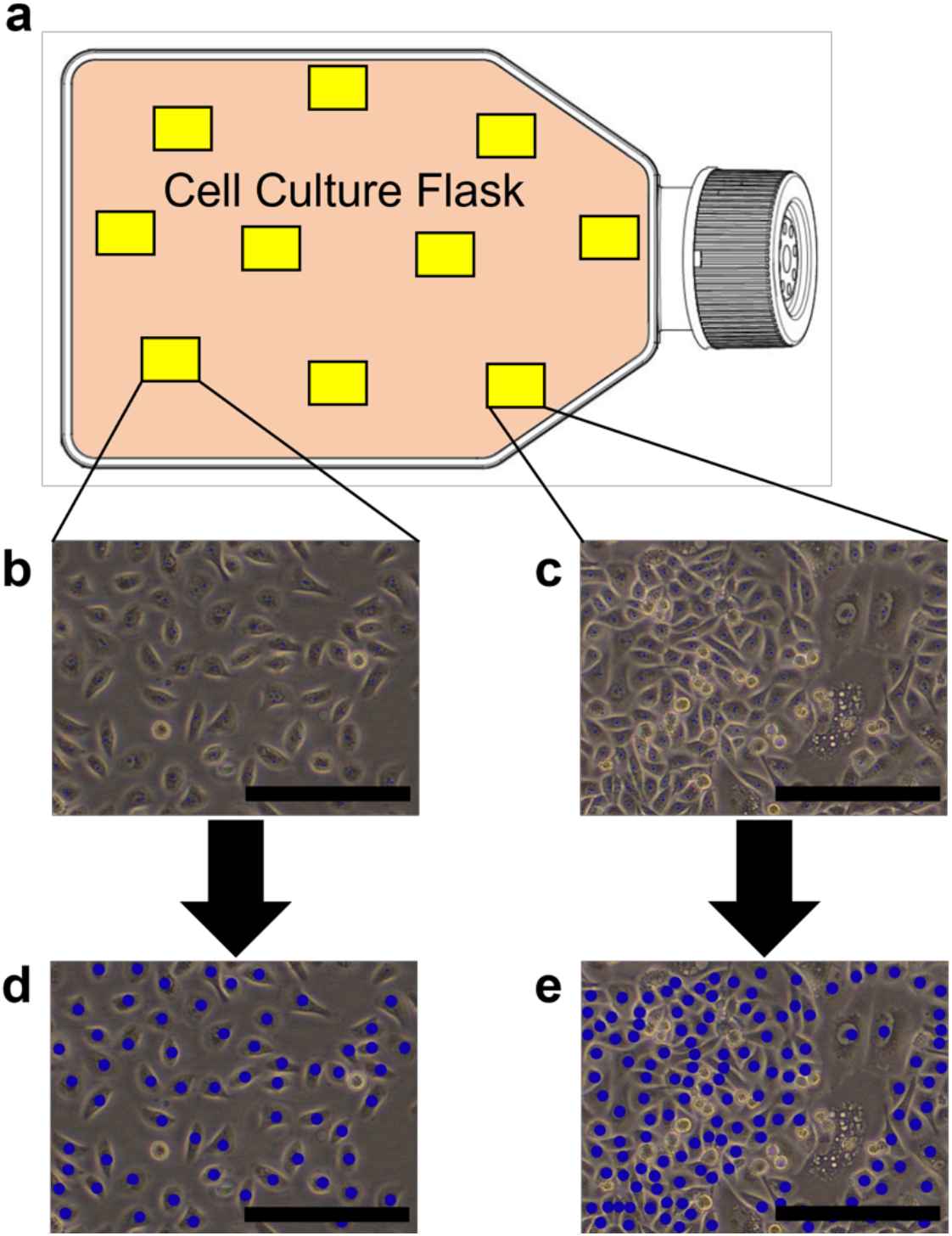
Depiction of the cell counting procedure. **(a)** Schematic of a T25 cell culture flask. Prior to conducting electrophysiology experiments, the flask for each cell line was imaged 10 times pseudo-randomly. Example imaging locations shown by yellow rectangles. **(b)** Image from the flask prior to counting is shown. Black scale bar indicates 200 μm. **(c)** An image from a different location within the same flask is shown. **(d, e)** The same images from **(b, c)** are shown post counting with all live cells marked by a blue dot using FIJI/ImageJ. The mean of the 10 images were taken to determine the total cell count of each flask.

**Figure S4:**
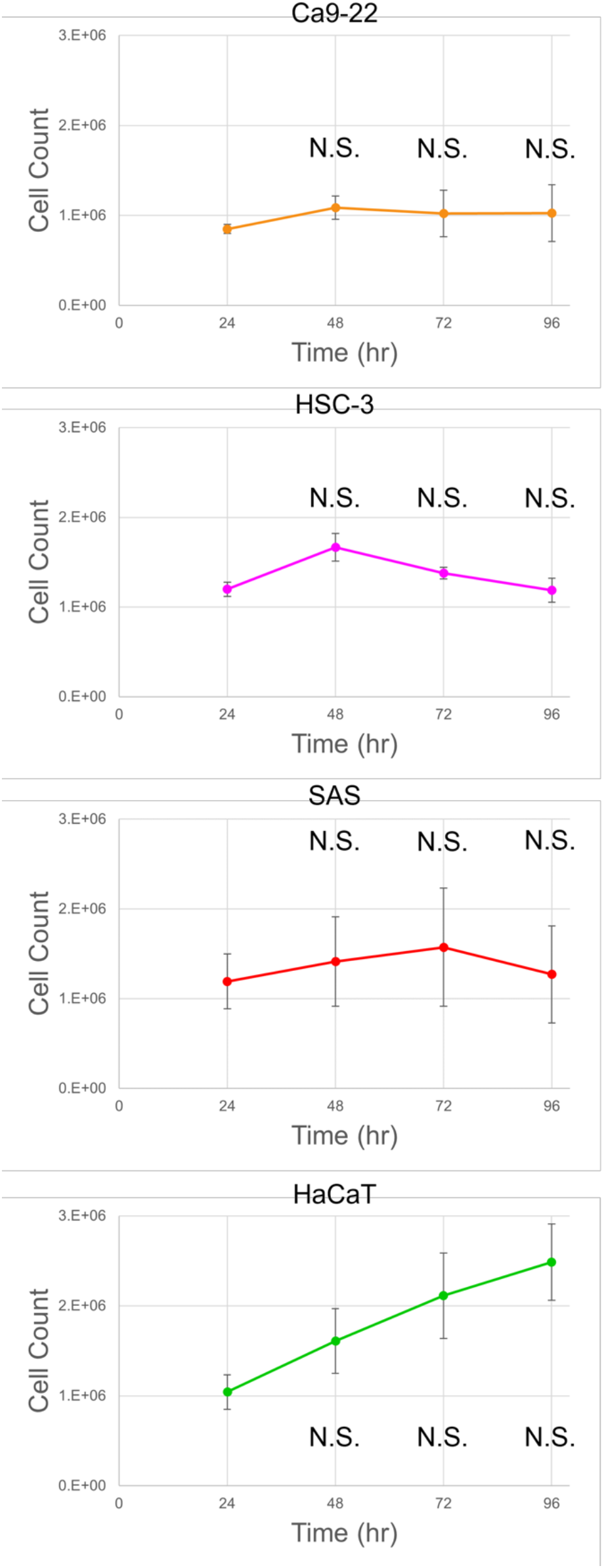
Cell growth curves of all four cell lines. Initially, cells were seeded at 1 x 10^6^ cells per flask at 0 h. Cells were counted from each flask at 24-hour intervals after seeding (24-, 48-, 72-, and 96-h). At each time point, the total cell count of each flask was averaged over seven replicates. Error bars are S.E.M from the seven replicates. No significant difference in cell counts were observed between 24-h and 48-, 72-, or 96-h (P < 0.05, d.f. = 6, 16, one-way ANOVA with Bonferroni correction).

**Figure S5:**
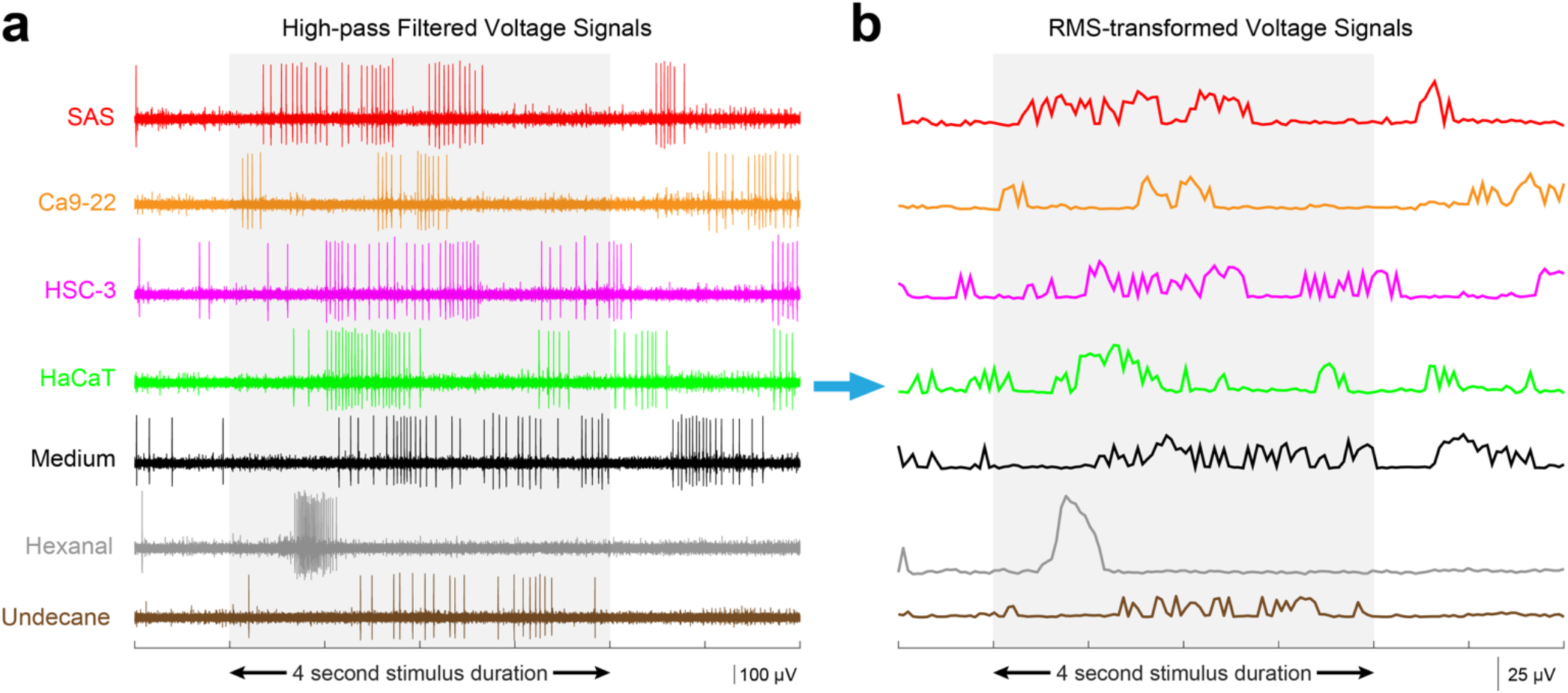
Root-mean squared (R.M.S) transformation largely preserved stimulus-specific spiking dynamics. **(a)** Representative recordings from an individual electrode are shown for all stimuli after high-pass filtering. The gray box delineates the 4 s stimulus presentation period. **(b)** R.M.S. transformed data traces of **panel a** recording (see Methods) reflect the spiking rate-based response dynamics, while reducing computational overhead. The gray box delineates the same 4 s stimulus presentation period as in **panel a**.

**Figure S6:**
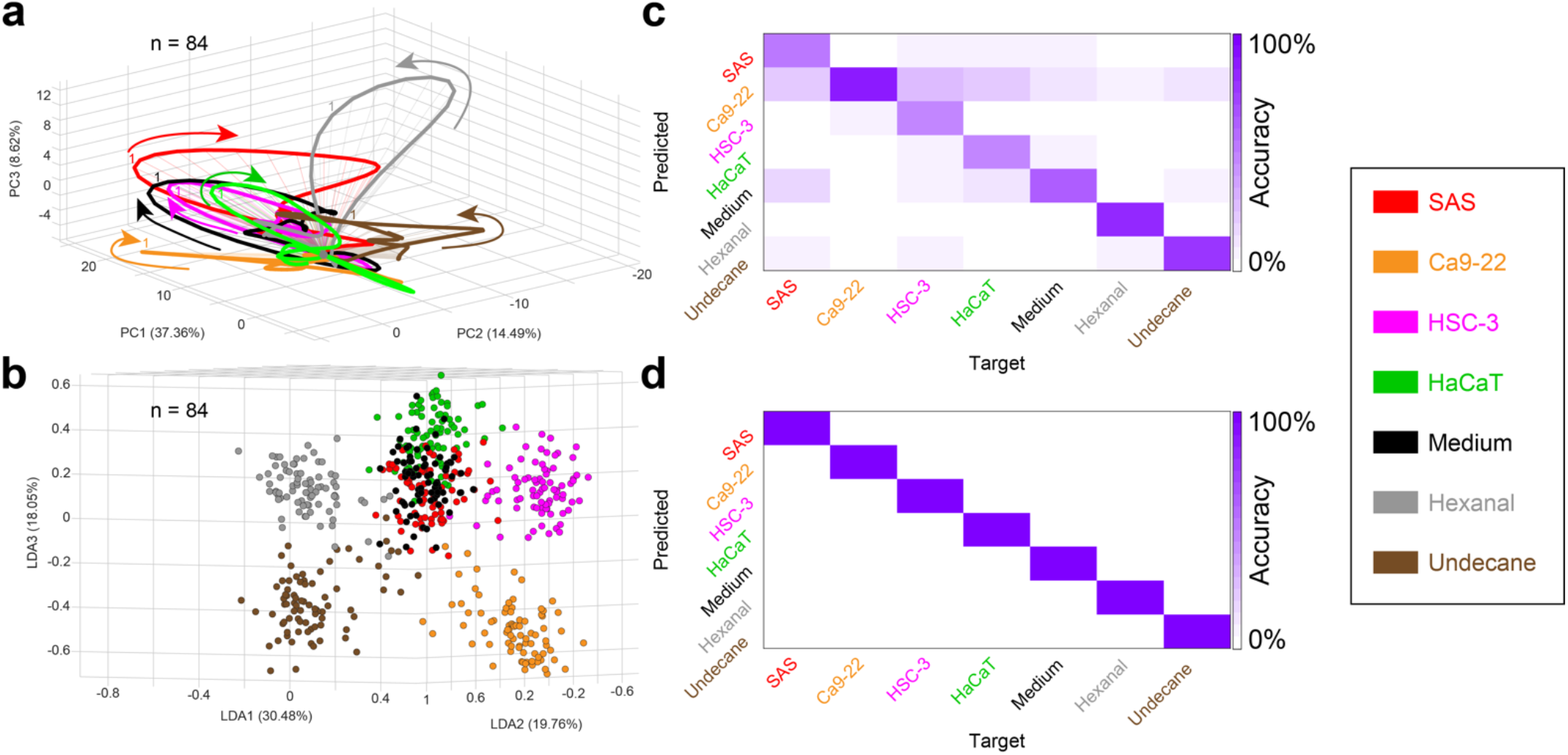
Using R.M.S transformed population PN voltage responses to classify the stimulus panel. Similar plots as shown in **Figure S1**, but here, we have used R.M.S transformed PN voltages to generate, **(a)** VOC evoked ensemble neural trajectories after PCA; **(b)** PN response clusters after LDA; **(c)** Bin-wise confusion matrix in high dimensional space; and **(d)** Trial-wise confusion matrix corresponding to all 7 stimuli. Spike-based and R.M.S-filtered PN time-series data both yielded excellent classification for all VOCs tested.

**Figure S7:**
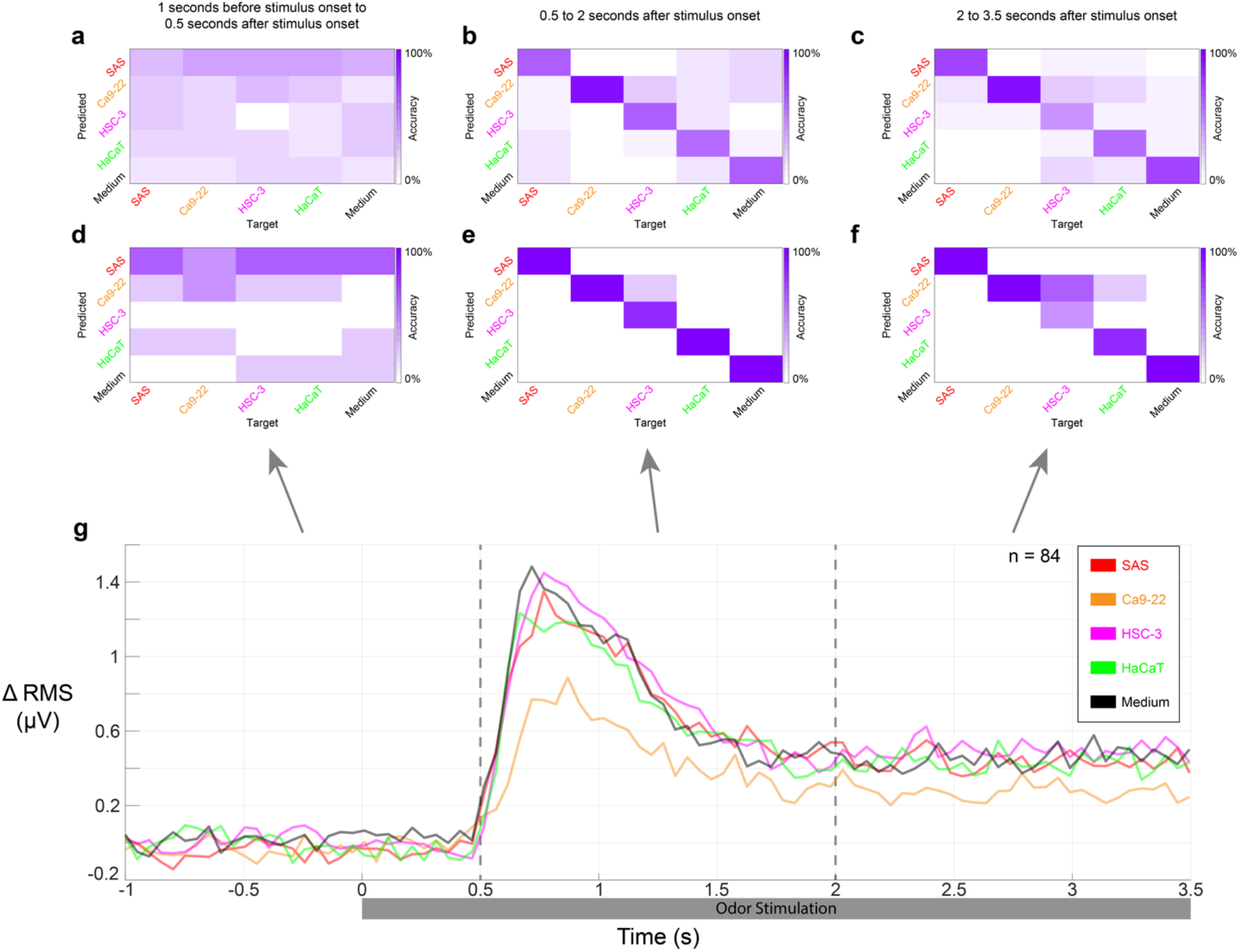
Cancer VOC classification during transient vs. steady state response periods of PN response. Time bin-wise confusion matrix analysis results shown for three 1.5 s duration time periods- **(a)** 1 s prior to 0.5 s after final olfactometer valve opening (pre-stimulus period). **b)** 0.5 to 2 s after final olfactometer valve opening (transient state), and **(c)** 2 to 3.5 s after final olfactometer valve opening (steady state). The pre-stimulus time period shows no stimulus-specific classification as expected because VOCs had not yet reached the antenna at this time. The transient state period (0.5 to 2 s) shows the best bin-wise classification of all 5 VOCs. The steady state period (2 to 3.5 s) also shows high classification success, but the diagonal values are relatively lower compared to the transient state period. **(d-f)** Trial-wise confusion matrices are shown for the same time windows as in **panel a**. **(g)** A population-based peri-stimulus time histogram (PSTH) plots the change in R.M.S. transformed values of all recording positions (*n* = 84) as a function of time. Time labels along the X-axis are relative to the stimulus onset time. A significant change in R.M.S. values is seen approximately 500 ms after the final valve was opened (i.e., stimulus onset).

## Contributions

D.S. conceptualized the study. D.S., A.F., M.P., E.H.A, and C.H.C designed experimental plans. A.F. and M.P. conducted electrophysiology experiments. M.P., E.H.A, and N.L. performed cell culture and imaging studies. C.H.C. provided resources and supervised the cell culture work. Data analysis was done by A.F., M.P, and D.S. Locust colony was maintained by E.C. Paper was written by D.S., A.F., and M.P. All the authors contributed to review and editing of the manuscript. D.S. performed overall project supervision and administration.

## Declaration of competing interests

The authors declare that they have no competing financial interests.

## Notes

### Competing Interest Statement

The authors have declared no competing interest.

## References

1 Pereira, J. et al. Breath analysis as a potential and non-invasive frontier in disease diagnosis: an overview. Metabolites 5, 3–55, doi:10.3390/metabo5010003 (2015).

2 Li, J. et al. Investigation of potential breath biomarkers for the early diagnosis of breast cancer using gas chromatography–mass spectrometry. Clinica Chimica Acta 436, 59–67, doi:https://doi.org/10.1016/j.cca.2014.04.030 (2014).

3 Miekisch, W., Schubert, J. K. & Noeldge-Schomburg, G. F. E. Diagnostic potential of breath analysis - Focus on volatile organic compounds. Clinica Chimica Acta 347, 25–39, doi:10.1016/j.cccn.2004.04.023 (2004).

4 Oakley-Girvan, I. & Davis, S. W. Breath based volatile organic compounds in the detection of breast, lung, and colorectal cancers: A systematic review. Cancer Biomark 21, 29–39, doi:10.3233/cbm-170177 (2017).

5 Phillips, M. et al. Prediction of breast cancer risk with volatile biomarkers in breath. Breast Cancer Res Treat 170, 343–350, doi:10.1007/s10549-018-4764-4 (2018).

6 Altomare, D. F. et al. Exhaled volatile organic compounds identify patients with colorectal cancer. Br J Surg 100, 144–150, doi:10.1002/bjs.8942 (2013).

7 Phillips, M. et al. Volatile organic compounds in breath as markers of lung cancer: a crosssectional study. Lancet 353, 1930–1933, doi:10.1016/s0140-6736(98)07552-7 (1999).

8 Kaloumenou, M., Skotadis, E., Lagopati, N., Efstathopoulos, E. & Tsoukalas, D. Breath Analysis: A Promising Tool for Disease Diagnosis-The Role of Sensors. Sensors (Basel) 22, doi:10.3390/s22031238 (2022).

9 Gouzerh, F. et al. Odors and cancer: Current status and future directions. Biochim Biophys Acta Rev Cancer 1877, 188644, doi:10.1016/j.bbcan.2021.188644 (2022).

10 Lüdke, A. & Galizia, C. G. Sniffing cancer: Will the fruit fly beat the dog? ChemoSense 15, 3–15 (2014).

11 Phillips, M. et al. Prediction of breast cancer using volatile biomarkers in the breath. Breast cancer research and treatment 99, 19–21 (2006).

12 Baldini, C. et al. Electronic Nose as a Novel Method for Diagnosing Cancer: A Systematic Review. Biosensors (Basel) 10, doi:10.3390/bios10080084 (2020).

13 Röck, F., Barsan, N. & Weimar, U. Electronic Nose: Current Status and Future Trends. Chemical Reviews 108, 705–725, doi:10.1021/cr068121q (2008).

14 Ko, H. J. & Park, T. H. Bioelectronic nose and its application to smell visualization. J Biol Eng 10, 17, doi:10.1186/s13036-016-0041-4 (2016).

15 DeGreeff, L. E. & Schultz, C. A. Canines: The Original Biosensors (1st ed.). 1st edn, (Jenny Stanford Publishing, 2022).

16 Ehmann, R. et al. Canine scent detection in the diagnosis of lung cancer: revisiting a puzzling phenomenon. European respiratory journal 39, 669–676 (2012).

17 Gordon, R. T. et al. The use of canines in the detection of human cancers. The Journal of Alternative and Complementary Medicine 14, 61–67 (2008).

18 Jezierski, T., Walczak, M., Ligor, T., Rudnicka, J. & Buszewski, B. Study of the art: canine olfaction used for cancer detection on the basis of breath odour. Perspectives and limitations. Journal of breath research 9, 027001 (2015).

19 McCulloch, M. et al. Diagnostic accuracy of canine scent detection in early-and late-stage lung and breast cancers. Integrative cancer therapies 5, 30–39 (2006).

20 Pickel, D., Manucy, G. P., Walker, D. B., Hall, S. B. & Walker, J. C. Evidence for canine olfactory detection of melanoma. Applied Animal Behaviour Science 89, 107–116 (2004).

21 Sonoda, H. et al. Colorectal cancer screening with odour material by canine scent detection. Gut 60, 814–819 (2011).

22 Saha, D. et al. Explosive sensing with insect-based biorobots. Biosensors and Bioelectronics: X 6, 100050, doi:https://doi.org/10.1016/j.biosx.2020.100050 (2020).

23 Stopfer, M., Jayaraman, V. & Laurent, G. Intensity versus Identity Coding in an Olfactory System. Neuron 39, 991–1004, doi:https://doi.org/10.1016/j.neuron.2003.08.011 (2003).

24 Saha, D. et al. A spatiotemporal coding mechanism for background-invariant odor recognition. Nature Neuroscience 16, 1830–1839, doi:10.1038/nn.3570 (2013).

25 Broome, B. M., Jayaraman, V. & Laurent, G. Encoding and Decoding of Overlapping Odor Sequences. Neuron 51, 467–482, doi:https://doi.org/10.1016/j.neuron.2006.07.018 (2006).

26 Nizampatnam, S., Saha, D., Chandak, R. & Raman, B. Dynamic contrast enhancement and flexible odor codes. Nature communications 9, 3062, doi:10.1038/s41467-018-05533-6 (2018).

27 Saha, D. et al. A spatiotemporal coding mechanism for background-invariant odor recognition. Nat Neurosci 16, 1830–1839, doi:10.1038/nn.3570 (2013).

28 Saha, D. et al. Behavioural correlates of combinatorial versus temporal features of odour codes. Nature communications 6, 6953, doi:10.1038/ncomms7953 (2015).

29 Saha, D. et al. Engaging and disengaging recurrent inhibition coincides with sensing and unsensing of a sensory stimulus. Nature communications 8, 15413, doi:10.1038/ncomms15413 (2017).

30 Bazhenov, M. et al. Model of cellular and network mechanisms for odor-evoked temporal patterning in the locust antennal lobe. Neuron 30, 569–581, doi:10.1016/s0896-6273(01)00286-0 (2001).

31 Bazhenov, M., Stopfer, M., Sejnowski, T. J. & Laurent, G. Fast odor learning improves reliability of odor responses in the locust antennal lobe. Neuron 46, 483–492, doi:S0896-6273(05)00278-3 [pii] 10.1016/j.neuron.2005.03.022 (2005).

32 Brown, S. L., Joseph, J. & Stopfer, M. Encoding a temporally structured stimulus with a temporally structured neural representation. Nature Neuroscience 8, 1568–1576 (2005).

33 Finelli, L. A., Haney, S., Bazhenov, M., Stopfer, M. & Sejnowski, T. J. Synaptic Learning Rules and Sparse Coding in a Model Sensory System. PLOS Computational Biology 4, e1000062, doi:10.1371/journal.pcbi.1000062 (2008).

34 Ito, I., Ong, R. C., Raman, B. & Stopfer, M. Sparse odor representation and olfactory learning. Nature Neuroscience 11, 1177–1184 (2008).

35 Stopfer, M., Bhagavan, S., Smith, B. H. & Laurent, G. Impaired odour discrimination on desynchronization of odour-encoding neural assemblies. Nature 390, 70–74 (1997).

36 Stopfer, M. & Laurent, G. Short-term memory in olfactory network dynamics. Nature 402, 664–668, doi:10.1038/45244 (1999).

37 Broome, B. M., Jayaraman, V. & Laurent, G. Encoding and decoding of overlapping odor sequences. Neuron 51, 467–482 (2006).

38 Cassenaer, S. & Laurent, G. Hebbian STDP in mushroom bodies facilitates the synchronous flow of olfactory information in locusts. Nature 448, 709–713, doi:10.1038/nature05973 (2007).

39 Cassenaer, S. & Laurent, G. Conditional modulation of spike-timing-dependent plasticity for olfactory learning. Nature 482, 47–52, doi:10.1038/nature10776 (2012).

40 Jortner, R. A., Farivar, S. S. & Laurent, G. A simple connectivity scheme for sparse coding in an olfactory system. The Journal of neuroscience: the official journal of the Society for Neuroscience 27, 1659–1669, doi:10.1523/JNEUROSCI.4171-06.2007 (2007).

41 Laurent, G. Dynamical representation of odors by oscillating and evolving neural assemblies. Trends Neurosci 19, 489–496, doi:10.1016/s0166-2236(96)10054-0 (1996).

42 Laurent, G. Olfactory network dynamics and the coding of multidimensional signals. Nature Reviews Neuroscience 3, 884–895, doi:10.1038/nrn964 (2002).

43 Laurent, G. & Davidowitz, H. Encoding of olfactory information with oscillating neural assemblies. Science 265, 1872–1875 (1994).

44 Papadopoulou, M., Cassenaer, S., Nowotny, T. & Laurent, G. Normalization for Sparse Encoding of Odors by a Wide-Field Interneuron. Science 332, 721, doi:10.1126/science.1201835 (2011).

45 Perez-Orive, J., Bazhenov, M. & Laurent, G. Intrinsic and circuit properties favor coincidence detection for decoding oscillatory input. The Journal of neuroscience: the official journal of the Society for Neuroscience 24, 6037–6047, doi:10.1523/jneurosci.1084-04.2004 (2004).

46 Perez-Orive, J. et al. Oscillations and Sparsening of Odor Representations in the Mushroom Body. Science 297, 359, doi:10.1126/science.1070502 (2002).

47 Turner, G. C., Bazhenov, M. & Laurent, G. Olfactory representations by Drosophila mushroom body neurons. Journal of neurophysiology 99, 734–746, doi:10.1152/jn.01283.2007 (2008).

48 Saha, D., Leong, K., Katta, N. & Raman, B. Multi-unit Recording Methods to Characterize Neural Activity in the Locust (Schistocerca Americana) Olfactory Circuits. Journal of visualized experiments: JoVE, doi: 10.3791/50139 (2013).

49 Vosshall, L. B., Amrein, H., Morozov, P. S., Rzhetsky, A. & Axel, R. A spatial map of olfactory receptor expression in the Drosophila antenna. Cell 96, 725–736, doi:10.1016/s0092-8674(00)80582-6 (1999).

50 Hallem, E. A. & Carlson, J. R. Coding of odors by a receptor repertoire. Cell 125, 143–160, doi:10.1016/j.cell.2006.01.050 (2006).

51 Mazor, O. & Laurent, G. Transient Dynamics versus Fixed Points in Odor Representations by Locust Antennal Lobe Projection Neurons. Neuron 48, 661–673, doi:https://doi.org/10.1016/j.neuron.2005.09.032 (2005).

52 Marumo, T. et al. Flavinated SDHA Underlies the Change in Intrinsic Optical Properties of Oral Cancers (2022 under review).

53 Pouzat, C., Mazor, O. & Laurent, G. Using noise signature to optimize spike-sorting and to assess neuronal classification quality. Journal of Neuroscience Methods 122, 43–57, doi:https://doi.org/10.1016/S0165-0270(02)00276-5 (2002).

54 Kharitonov, S. A. & Barnes, P. J. Exhaled markers of pulmonary disease. American journal of respiratory and critical care medicine 163, 1693–1722 (2001).

55 Paredi, P. et al. Exhaled ethane is elevated in cystic fibrosis and correlates with carbon monoxide levels and airway obstruction. American journal of respiratory and critical care medicine 161, 1247–1251 (2000).

56 Barker, M. et al. Volatile organic compounds in the exhaled breath of young patients with cystic fibrosis. European Respiratory Journal 27, 929–936 (2006).

57 Paredi, P., Kharitonov, S. A. & Barnes, P. J. Elevation of exhaled ethane concentration in asthma. American journal of respiratory and critical care medicine 162, 1450–1454 (2000).

58 Paredi, P. et al. Exhaled ethane, a marker of lipid peroxidation, is elevated in chronic obstructive pulmonary disease. American journal of respiratory and critical care medicine 162, 369–373 (2000).

59 Phillips, M. et al. Volatile organic compounds in breath as markers of lung cancer: a crosssectional study. The Lancet 353, 1930–1933 (1999).

60 Phillips, M. et al. Volatile markers of breast cancer in the breath. The breast journal 9, 184–191 (2003).

61 Cazzola, M. et al. Outcomes for COPD pharmacological trials: from lung function to biomarkers. European respiratory journal 31, 416–469 (2008).

62 De Heer, K. et al. Detection of airway colonization by Aspergillus fumigatus by use of electronic nose technology in patients with cystic fibrosis. Journal of clinical microbiology 54, 569–575 (2016).

63 Bikov, A. et al. Expiratory flow rate, breath hold and anatomic dead space influence electronic nose ability to detect lung cancer. BMC pulmonary medicine 14, 1–9 (2014).

64 Kahn, N., Lavie, O., Paz, M., Segev, Y. & Haick, H. Dynamic nanoparticle-based flexible sensors: diagnosis of ovarian carcinoma from exhaled breath. Nano letters 15, 7023–7028 (2015).

65 Behera, B., Joshi, R., Vishnu, G. K. A., Bhalerao, S. & Pandya, H. J. Electronic nose: A non-invasive technology for breath analysis of diabetes and lung cancer patients. Journal of breath research 13, 024001 (2019).

66 De Vries, R. et al. Integration of electronic nose technology with spirometry: validation of a new approach for exhaled breath analysis. Journal of breath research 9, 046001 (2015).

67 Bach, J.-P. et al. Measuring compounds in exhaled air to detect Alzheimer’s disease and Parkinson’s disease. PloS one 10, e0132227 (2015).

68 Kunos, L. et al. Evening and morning exhaled volatile compound patterns are different in obstructive sleep apnoea assessed with electronic nose. Sleep and Breathing 19, 247–253 (2015).

69 Leal, R. V. et al. in Proceedings of the 2019 9th International Conference on Biomedical Engineering and Technology. 13–17 (2019).

70 Sethi, S., Nanda, R. & Chakraborty, T. Clinical application of volatile organic compound analysis for detecting infectious diseases. Clinical microbiology reviews 26, 462–475 (2013).

71 Sun, X., Shao, K. & Wang, T. Detection of volatile organic compounds (VOCs) from exhaled breath as noninvasive methods for cancer diagnosis. Analytical and bioanalytical chemistry 408, 2759–2780 (2016).

72 Martin, A. N., Farquar, G. R., Jones, A. D. & Frank, M. Human breath analysis: methods for sample collection and reduction of localized background effects. Analytical and bioanalytical chemistry 396, 739–750 (2010).

73 O’Neill, H. J., Gordon, S. M., O’Neill, M. H., Gibbons, R. D. & Szidon, J. P. A computerized classification technique for screening for the presence of breath biomarkers in lung cancer. Clinical Chemistry 34, 1613–1618, doi:10.1093/clinchem/34.8.1613 (1988).

74 Sancho, P. et al. MYC/PGC-1α Balance Determines the Metabolic Phenotype and Plasticity of Pancreatic Cancer Stem Cells. Cell Metab 22, 590–605, doi:10.1016/j.cmet.2015.08.015 (2015).

75 Williams, H. & Pembroke, A. Sniffer dogs in the melanoma clinic? The Lancet 333, 734 (1989).

76 Willis, C. M. et al. Olfactory detection of human bladder cancer by dogs: proof of principle study. Bmj 329, 712 (2004).

77 Urbanová, L., Vyhnánková, V., Krisová, Š., Pacík, D. & Nečas, A. Intensive training technique utilizing the dog’s olfactory abilities to diagnose prostate cancer in men. Acta Veterinaria Brno 84 (2015).

78 Cornu, J.-N., Cancel-Tassin, G., Ondet, V., Girardet, C. & Cussenot, O. Olfactory detection of prostate cancer by dogs sniffing urine: a step forward in early diagnosis. European urology 59, 197–201 (2011).

79 Horvath, G., Andersson, H. & Paulsson, G. Characteristic odour in the blood reveals ovarian carcinoma. EUROMEDICA, 67 (2011).

80 Los, E. A., Ramsey, K. L., Guttmann-Bauman, I. & Ahmann, A. J. Reliability of trained dogs to alert to hypoglycemia in patients with type 1 diabetes. Journal of diabetes science and technology 11, 506–512 (2017).

81 Bomers, M. K. et al. Using a dog’s superior olfactory sensitivity to identify Clostridium difficile in stools and patients: proof of principle study. Bmj 345 (2012).

82 Eskandari, E. et al. Sniffer dogs as a screening/diagnostic tool for COVID-19: a proof of concept study. BMC infectious diseases 21, 1–8 (2021).

83 Mgode, G. F. et al. Diagnosis of tuberculosis by trained African giant pouched rats and confounding impact of pathogens and microflora of the respiratory tract. Journal of clinical microbiology 50, 274–280 (2012).

84 Weetjens, B. et al. African pouched rats for the detection of pulmonary tuberculosis in sputum samples. The International journal of tuberculosis and lung disease 13, 737–743 (2009).

85 Kontos, E. et al. Bees can be trained to identify SARS-CoV-2 infected samples. bioRxiv (2021).

86 Suckling, D. M. & Sagar, R. L. Honeybees Apis mellifera can detect the scent of Mycobacterium tuberculosis. Tuberculosis 91, 327–328 (2011).

87 Strauch, M. et al. More than apples and oranges-Detecting cancer with a fruit fly’s antenna. Scientific reports 4, 1–9 (2014).

88 Piqueret, B. et al. Ants detect cancer cells through volatile organic compounds. Iscience 25, 103959 (2022).

89 Cheema, J. A., Carraher, C., Plank, N. O. V., Travas-Sejdic, J. & Kralicek, A. Insect odorant receptor-based biosensors: Current status and prospects. Biotechnology Advances 53, 107840, doi:https://doi.org/10.1016/j.biotechadv.2021.107840 (2021).

90 Feng, X. et al. A living cell-based biosensor utilizing G-protein coupled receptors: principles and detection methods. Biosens Bioelectron 22, 3230–3237, doi:10.1016/j.bios.2007.03.002 (2007).

91 Lee, S. H., Oh, E. H. & Park, T. H. Cell-based microfluidic platform for mimicking human olfactory system. Biosens Bioelectron 74, 554–561, doi:10.1016/j.bios.2015.06.072 (2015).

92 Hwi Jin, K. & Tai Hyun, P. Dual signal transduction mediated by a single type of olfactory receptor expressed in a heterologous system. Biological Chemistry 387, 59–68, doi:https://doi.org/10.1515/BC.2006.009 (2006).

93 Lee, S. H., Jun, S. B., Ko, H. J., Kim, S. J. & Park, T. H. Cell-based olfactory biosensor using microfabricated planar electrode. Biosensors and Bioelectronics 24, 2659–2664, doi:https://doi.org/10.1016/j.bios.2009.01.035 (2009).

94 Lee, S. H., Jeong, S. H., Jun, S. B., Kim, S. J. & Park, T. H. Enhancement of cellular olfactory signal by electrical stimulation. ELECTROPHORESIS 30, 3283–3288, doi:10.1002/elps.200900124 (2009).

95 Misawa, N. et al. Construction of a Biohybrid Odorant Sensor Using Biological Olfactory Receptors Embedded into Bilayer Lipid Membrane on a Chip. ACS Sens 4, 711–716, doi:10.1021/acssensors.8b01615 (2019).

96 Mitsuno, H., Sakurai, T., Namiki, S., Mitsuhashi, H. & Kanzaki, R. Novel cell-based odorant sensor elements based on insect odorant receptors. Biosens Bioelectron 65, 287–294, doi:10.1016/j.bios.2014.10.026 (2015).

97 Termtanasombat, M. et al. Cell-Based Odorant Sensor Array for Odor Discrimination Based on Insect Odorant Receptors. J Chem Ecol 42, 716–724, doi:10.1007/s10886-016-0726-7 (2016).

98 Murugathas, T. et al. Biosensing with Insect Odorant Receptor Nanodiscs and Carbon Nanotube Field-Effect Transistors. ACS Appl Mater Interfaces 11, 9530–9538, doi:10.1021/acsami.8b19433 (2019).

99 Sato, K. & Takeuchi, S. Chemical Vapor Detection Using a Reconstituted Insect Olfactory Receptor Complex. Angewandte Chemie International Edition 53, 11798–11802, doi:10.1002/anie.201404720 (2014).

100 Boukamp, P. et al. Normal keratinization in a spontaneously immortalized aneuploid human keratinocyte cell line. Journal of Cell Biology 106, 761–771, doi:10.1083/jcb.106.3.761 (1988).

101 Chen, P. et al. A Novel Peptide for Simultaneously Enhanced Treatment of Head and Neck Cancer and Mitigation of Oral Mucositis. PLOS ONE 11, e0152995, doi:10.1371/journal.pone.0152995 (2016).

